# Lipocalins are versatile regulators of development and stress response in mosses

**DOI:** 10.1101/2025.03.13.638958

**Authors:** Shuanghua Wang, Jianchao Ma, Qia Wang, Yanlong Guan, Xiangyang Hu, Hang Sun, Jinling Huang

**Affiliations:** Key Laboratory for Plant Diversity and Biogeography of East Asia, Yunnan Key Laboratory for Fungal Diversity and Green Development, Kunming Institute of Botany, Chinese Academy of Sciences, Kunming, 650201, China; State Key Laboratory of Crop Stress Adaptation and Improvement, Key Laboratory of Plant Stress Biology, School of Life Sciences, Henan University, Kaifeng, 475001, China; School of Life Sciences, Shanghai University, Shanghai, 200444, China; Department of Biology, East Carolina University, Greenville, NC, USA

## Abstract

Major physiological and developmental innovations, such as stress tolerance strategies and three-dimensional growth, are key to land plant adaptation and evolution, but their genetic basis remains to be fully understood. Although lipocalins are known for their role to optimize adaptation and development in animals, they are heavily understudied in plants. Here we show that lipocalins in the moss *Physcomitrium patens* are versatile regulators of miscellaneous key physiological and developmental processes. Phylogenetics analyses revealed three major groups of plant lipocalins, including a newly identified group that is restricted to seedless plants. The temperature-induced lipocalin gene in *P. patens* (*PpTIL*) is functionally conserved with flowering plants in response to various abiotic stresses (e.g., heat, cold, salt, and dehydration). We demonstrate that *PpTIL* not only regulates protonemal development and the transition from two-dimensional to three-dimensional growth, but also affects many other processes such as lipid transport and metabolism, auxin biosynthesis and transport, and chlorophyll catabolism. These findings provide major insights into the role of lipocalins in land plant evolution.

## Introduction

The transition from water to land represents one of the most significant events in the evolution of plants^1–3^. To overcome the challenges in harsh terrestrial environments (e.g., water deficiency, strong ultraviolet radiation, surface temperature fluctuation, and pathogen infection)^2,4^, land plants evolved various physiological and developmental innovations, such as heat and dehydration tolerance, three-dimensional growth, and alternation of generations between gametophytes and sporophytes^5,6^. Bryophytes (liverworts, hornworts, and mosses) split from vascular plants early in land plant evolution^7–9^, and they are often thought to have retained many features of early land plants^6,10^. For instance, both two-dimensional (2D) and three-dimensional (3D) growth forms now co-exist during gametophyte development in the moss *Physcomitrium patens*^6,11,12^. Knowledge about the genetic basis of these innovations is key to understanding the origin and evolution of land plants^6,13^.

Lipocalins are a large, ancient and versatile family of proteins that are widely distributed in both prokaryotes and eukaryotes^14^. Members of the lipocalin family, although often varying considerably at sequence level, share a conserved β-barrel composed of eight antiparallel strands. The barrel is closed on one end and open on the opposite end, forming a calyx-like structure^15,16^. By binding small, mostly hydrophobic, molecules, cell-surface receptors, or forming complexes with soluble macromolecules, they fulfill numerous functions^15^. In bacteria, lipocalins are typically anchored to the outer membrane, where they are involved in resistance to antibiotics, starvation, salt, and other stresses^17,18^. In animals, lipocalins have been extensively studied. Particularly, the transition of vertebrates from water to land and their subsequent diversification were associated with rapid expansion and neofunctionalization of the lipocalin genes^19–21^. These lipocalin genes participate in ligand binding and transport, immune and defensive responses, chemical signaling, reproduction and development, and many other processes^15,22,23^. Plant lipocalins are structurally conserved with bacterial lipocalins, mammalian apolipoprotein D (ApoD), and insect Lazarillo, despite their usually modest protein sequence similarity (<31% identity)^24^. Within plants, lipocalins have been classified into two major groups, including chloroplastic lipocalins (CHLs) and temperature induced lipocalins (TILs), based on their sequence similarities and gene structures^25^. CHLs and TILs in flowering plants are localized in different cellular compartments, mostly known for their distinct but overlapping functions in protecting membrane lipids from reactive oxygen species (ROS) damages^26–29^. While CHLs are found within chloroplast thylakoid lumen and involved in light stress response^27,30^, TILs are typically localized at the plasma membrane and confer tolerance to other abiotic stresses (cold, heat, and salt)^26,29,31,32^. Recently, it has also been reported that the TIL protein in *Arabidopsis thaliana*, by interacting with retinal, regulates root clock oscillation and lateral root initiation^33^.

Despite their role in stress adaptation and root development in flowering plants, lipocalins have not been studied in bryophytes. Given the fact that lipocalins are involved in numerous processes of animal development and evolution, particularly the transition of vertebrates from water to land and ensuing diversification^20,21^, it is intriguing whether they played a similar role in plant colonization of land. In this study, we provide a comprehensive evolutionary analysis of plant lipocalins, including the identification of a major lipocalin group that is specific to seedless land plants and their charophyte algal relatives. Experimental evidence indicates that the *TIL* gene in the moss *P. patens* (*PpTIL*) is functionally conserved with flowering plants in response to abiotic stresses. Importantly, we show that *PpTIL* regulates a variety of other key physiological and developmental processes, including auxin biosynthesis and transport, transition from chloronema to caulonema, and three-dimensional development. These data suggest that lipocalins are versatile regulators in bryophytes and they likely played a pivotal role in plant colonization of land and subsequent diversification.

## Results

### Evolution of plant lipocalins

The genome of *P. patens* includes a single annotated gene copy of *TIL* (*PpTIL;* Phytozome ID *Pp3c18_22450*) and *CHL* (*PpCHL*; *Pp3c9_6110*), respectively. The *PpTIL* gene encodes a protein of 189 amino acids (aa) that is composed of a lipocalin-Blc-like (cd19438) or lipocalin_2 domain (PF08212) (Fig. 1a). The PpCHL protein is 341 aa in length with a lipocalin-CHL domain (cd19851) in the middle (Fig. 1a). We performed comprehensive BLAST searches of the NCBI non-redundant (nr) protein sequences and other related datasets, using both PpTIL and PpCHL protein sequences as query, as well as HMMER searches using the lipocalin and lipocalin-like domains (PF00061 and PF08212) (see Materials and Methods). Hits generated from the searches were distributed in streptophytes (land plants and charophyte algae), bacteria, viruses, animals, and miscellaneous other eukaryotes. In addition, we also found lipocalins in various archaeal groups, despite the common belief that lipocalins are absent in archaea^20^. Notably, plant TIL protein sequences are highly similar to their bacterial homologs (Fig. 1b; Supplementary Fig. 1), with a percent identity and coverage of up to 50.31% and 82%, respectively.

**Fig. 1.**
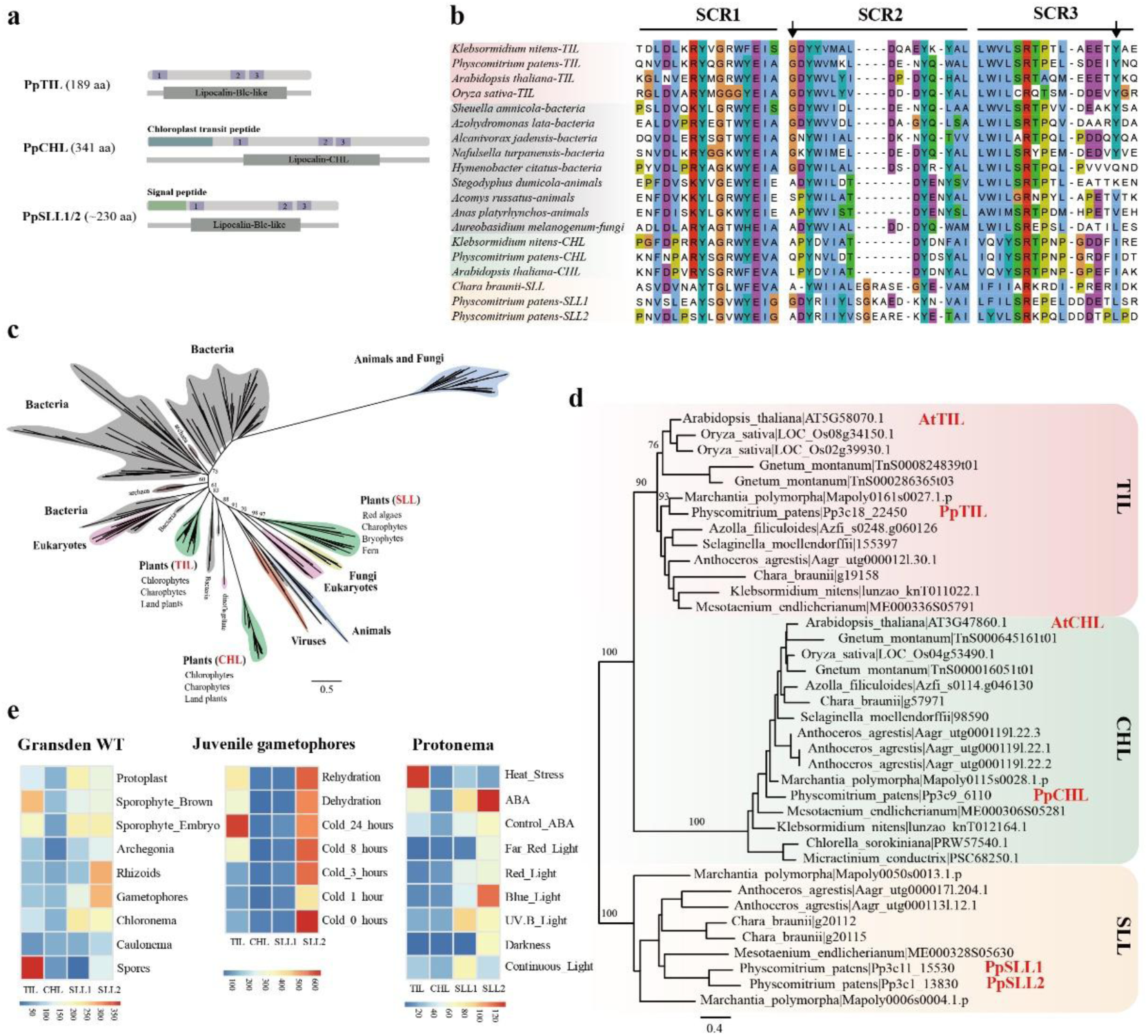
Evolution, signal prediction, and expression of plant lipocalins. **a,** Schematic representation of plant lipocalins. Lilac boxes labeled 1, 2, and 3 represent three structurally conserved regions (SCRs). Signal peptide of PpSLL and chloroplast transit peptide of PpCHL are marked in cyan and green, respectively. **b**, Three SCRs of lipocalin proteins from different groups. Arrows show relatively conserved amino acids between plant TIL and bacteria lipocalin proteins. **c,** Molecular phylogeny of lipocalin proteins sampled from all three domains of life (bacteria, archaea, and eukaryotes) and viruses. **d,** Molecular phylogeny of plant lipocalins. Numbers associated with main branches show ultrafast bootstrap values from maximum likelihood analyses in **c** and **d**. **e,** Expression heatmaps of four lipocalin genes in *P. patens*. Expression data were downloaded from the PEATmoss database (https://peatmoss.plantcode.cup.uni-freiburg.de/).

Phylogenetic analyses revealed three distinct groups of lipocalins in plants (Fig. 1c, d). Besides TILs and CHLs reported earlier^25^, a third major lipocalin group was identified in our analyses. Like TILs, members of this newly identified lipocalin group mostly consist of a lipocalin-Blc-like domain (Fig. 1a). In green plants, this group is largely restricted to charophytes and bryophytes but rarely found in vascular plants. We therefore refer to this newly identified group as seedless plants-specific lipocalins (SLLs). Two *SLL* genes are annotated in the genome of *P. patens* (*Pp3c11_15530* and *Pp3c1_13830*), each of which encodes a protein that is predicted by SignalP^34^ to contain a Sec/SPI signal peptide or by DeepLoc^35^ to be extracellular (Fig. 1a). Consistent with the results of sequence comparisons (Fig. 1b), TILs of green plants formed a strongly supported monophyletic group, which in turn was nested within bacterial sequences and appeared to be distantly related to other eukaryotic lipocalins (Fig. 1c). Based on the information from PEATmoss^36^, the three groups of plant lipocalins have distinct expression profiles in the lifecycle of *P. patens* or under various abiotic stresses (Fig. 1e; Supplementary Fig. 2).

### PpTIL is localized both in chloroplasts and at the plasma membrane

Because the *TIL* gene has been studied in flowering plants, we decided to perform comparative investigations of this gene in *P. patens*. The TIL protein in flowering plants is known to be localized at the plasma membrane^25,26,28^ and might migrate to symplast under salt stress^32^. To assess the subcellular localization of PpTIL, we generated *PpTIL*pro*:PpTIL-EGFP-GUS* transgenic plants by expressing the coding region of PpTIL fused with the EGFP and β-glucuronidase (GUS) reporter genes, under the control of the *PpTIL* native promoter (Supplementary Fig. 3). To our surprise, we observed GFP signal associated with both chloroplasts and the plasma membrane (Fig. 2a-d). In particular, the GFP intensity is significantly stronger in chloroplasts than at the plasma membrane.

**Fig. 2.**
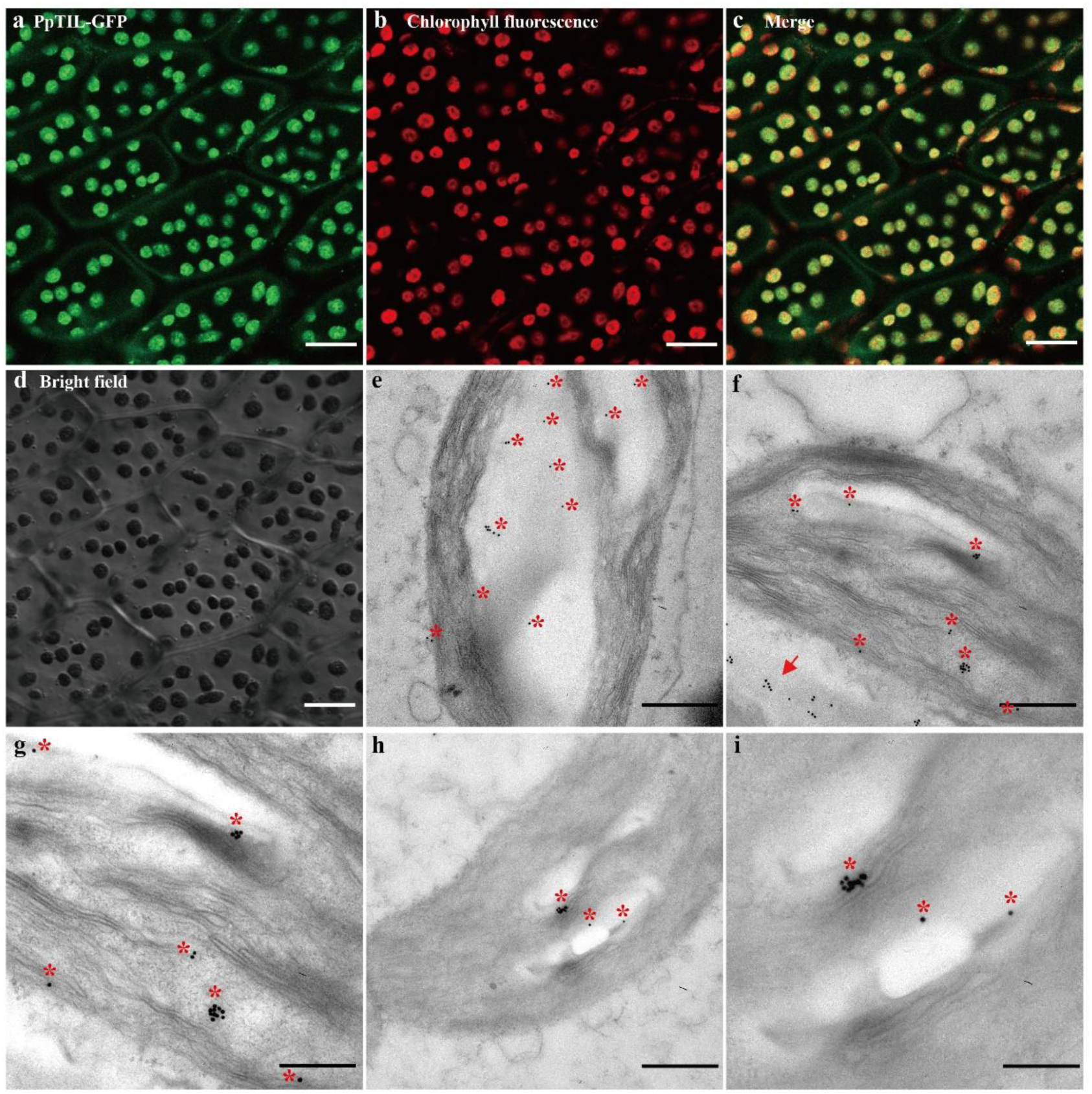
Subcellular localization of PpTIL in *P. patens.* **a,** GFP fluorescence of PpTIL. **b,** Chlorophyll fluorescence of chloroplast in **a**. **c,** Merge of **a** and **b**. **d,** Bright field of **a**, **b** and **c**. GFP signal of *PpTIL*pro*:PpTIL-EGFP-GUS* lines was observed in chloroplasts and at the plasma membrane. Micrograph images shown were observed from at least three biological replicates. Scale bar: 25 μm. **e-i,** Immunoelectron microscopy (IEM) of PpTIL. Colloidal gold nanoparticles (10 nm) are mainly located in chloroplasts stroma and thylakoid (red asterisks), and the plasma membrane (red arrow). **g** and **i** show enlargement of **f** and **h**, respectively. Scale bar: 500 nm in **e**, **f** and **h**, and 200 nm in **g** and **i**.

Because PpTIL lacks a transit peptide for chloroplast targeting (Fig. 1a), we further investigated whether PpTIL is indeed localized in chloroplasts and, if so, whether it is associated with stroma or thylakoid. We carried out immunoelectron microscopy (IEM) using protonemata and gametophores of *PpTIL*pro*:PpTIL-EGFP-GUS* plants, of which ultra-thin slices were prepared and incubated with anti-GFP antibody and colloidal gold-conjugated goat anti-rabbit IgG. We observed colloidal gold nanoparticles (10 nm) mainly in stroma and thylakoid of chloroplasts, as well as at the plasma membrane (Fig. 2e-i). These findings confirm the localization of PpTIL protein both in chloroplasts and at the plasma membrane.

### *PpTIL* is functionally conserved in abiotic stress response

In flowering plants, TILs are involved in tolerance of extreme temperatures, salt, and oxidative stresses^26,28,29,32^. Therefore, we first performed experiments to assess whether they have similar functions in bryophytes. Protonemata of *P. patens* were treated with heat, cold, salt, dehydration and ABA, respectively, for various durations, and the expression profile of *PpTIL* was assessed. We found that *PpTIL* expression was significantly up-regulated under these treatments and then gradually decreased to varying degrees (Fig. 3a-f), indicating that *PpTIL* is responsive to different abiotic stresses.

**Fig. 3.**
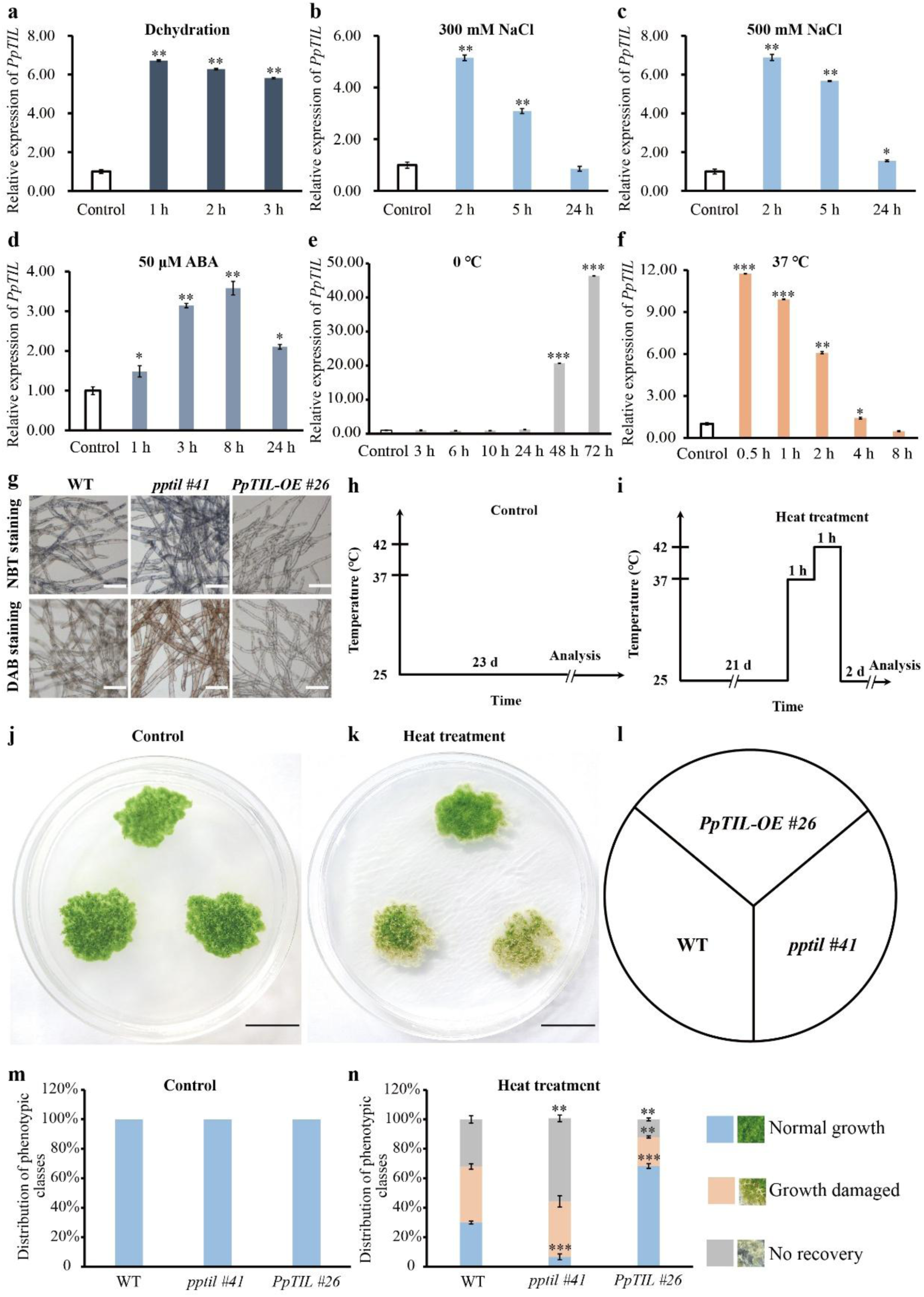
*PpTIL* is responsive to multiple abiotic stresses. **a-f,** Expression levels of *PpTIL* under control conditions and different abiotic stress treatments. *PpEF1a* was used as reference gene for normalization. Each bar shows mean ± s.e.m. of three biological replicates. **g,** ROS level revealed by DAB and NBT staining in protonemata of WT, *ko* and *OE* plants. **h,** Assay condition for thermotolerance under control condition. **i,** Assay condition for thermotolerance under heat treatment. d stands for days in **h** and **i**. **j,** Photographs of WT, *ko*, *OE* plants under control condition. **k,** Photographs of WT, *ko*, *OE* plants under heat treatment. Scale bar: 2 cm in **j** and **k**. WT, *ko* and *OE* plants in each treatment group were grown on the same plate, as shown in **l. m,** Quantification of the data shown in **j**. **n,** Quantification of the data shown in **k**. Asterisks indicate a statistically significant difference compared with the wild type based on a two-tailed Student’s *t* test (* *p*< 0.05, ** *p*< 0.01, *** *p*< 0.001).

We performed further detailed experiments to understand the role of *PpTIL* in heat stress tolerance. Transformation vectors for *PpTIL* gene knockout (*ko*) and overexpression (*OE*) were constructed, and four *ko* and three *OE* transformants were generated using long fragment homologous recombination and PEG-mediated protoplast transformation^37^. The *ko* and *OE* plants were identified by genomic PCR and real-time quantitative RT-PCR (qRT-PCR) (Supplementary Figs. 4-5). Additional chromosome ploidy analyses by flow cytometry showed that the ploidy of all generated *ko* and *OE* lines was consistent with that of the wild type (Supplementary Fig. 6). Three-week-old protonemata of WT, *ko* and *OE* plants grown on BCDAT medium were treated at 37℃ for one hour, followed by another hour at 42℃, and then recovered at 25℃ under a 16-hour light and 8-hour dark regime (Fig. 3h, i). We found that, compared to the wild type, the *ko* mutants were significantly more sensitive to heat stress whereas the *OE* lines were significantly more resistant (Fig. 3j-n). Specifically, only about 7% of *ko* mutants showed normal growth two days after the heat treatment, whereas 37% and 56% of them exhibited stunted growth or failed to recover from heat stress, respectively (Fig. 3k, n). In comparison, approximately 30% of WT plants grew normally, whereas 38% had stunted growth and 32% were unable to recover from the heat stress. Most *OE* plants (68%) exhibited normal growth, and only a small fraction showed retarded growth or failed to recover (20% and 12%, respectively) (Fig. 3k, n). These data further corroborate the conserved function of the *PpTIL* gene in thermotolerance.

Because TILs are known to protect plants and other organisms from oxidative stresses^29,31,38^, we also carried out diaminobenzidine (DAB) and nitroblue tetrazolium (NBT) staining to assess the reactive oxygen species (ROS) level in WT, *ko* and *OE* plants. Compared to the wild type, the ROS level was visibly higher in *ko* plants but lower in *OE* lines (Fig. 3g). These findings again suggest that *PpTIL* is functionally conserved in plant response to abiotic stresses and ROS reduction.

### *PpTIL* influences gametophore development in *P. patens*

In *P. patens*, the lifecycle is gametophyte dominant. Spores first germinate and develop into two-dimensional chloronemata, which in turn produce caulonemata through continuous division of tip-growing caulonemal initiation cells^12,39^. About 5% of the caulonemal cells lead to three-dimensional growth by changing the direction of the cell division plate^11,40^. Eventually, three-dimensional buds and gametophores formed. To assess the expression of *PpTIL* at different developmental stages of *P. patens*, we observed GFP and GUS signals using *PpTIL*pro*:PpTIL-EGFP-GUS* transgenic plants. GFP and GUS signals were detected in chloronemata, caulonemata, three-faced buds, leafy gametophores, archegonia and antheridia, and various other structures (Fig. 4a-j; Supplementary Fig. 7). Particularly, strong signal was observed in buds and apical meristems of gametophores (Fig. 4a-d, f-i), suggesting that *PpTIL* is involved in gametophore development in *P. patens*.

**Fig. 4.**
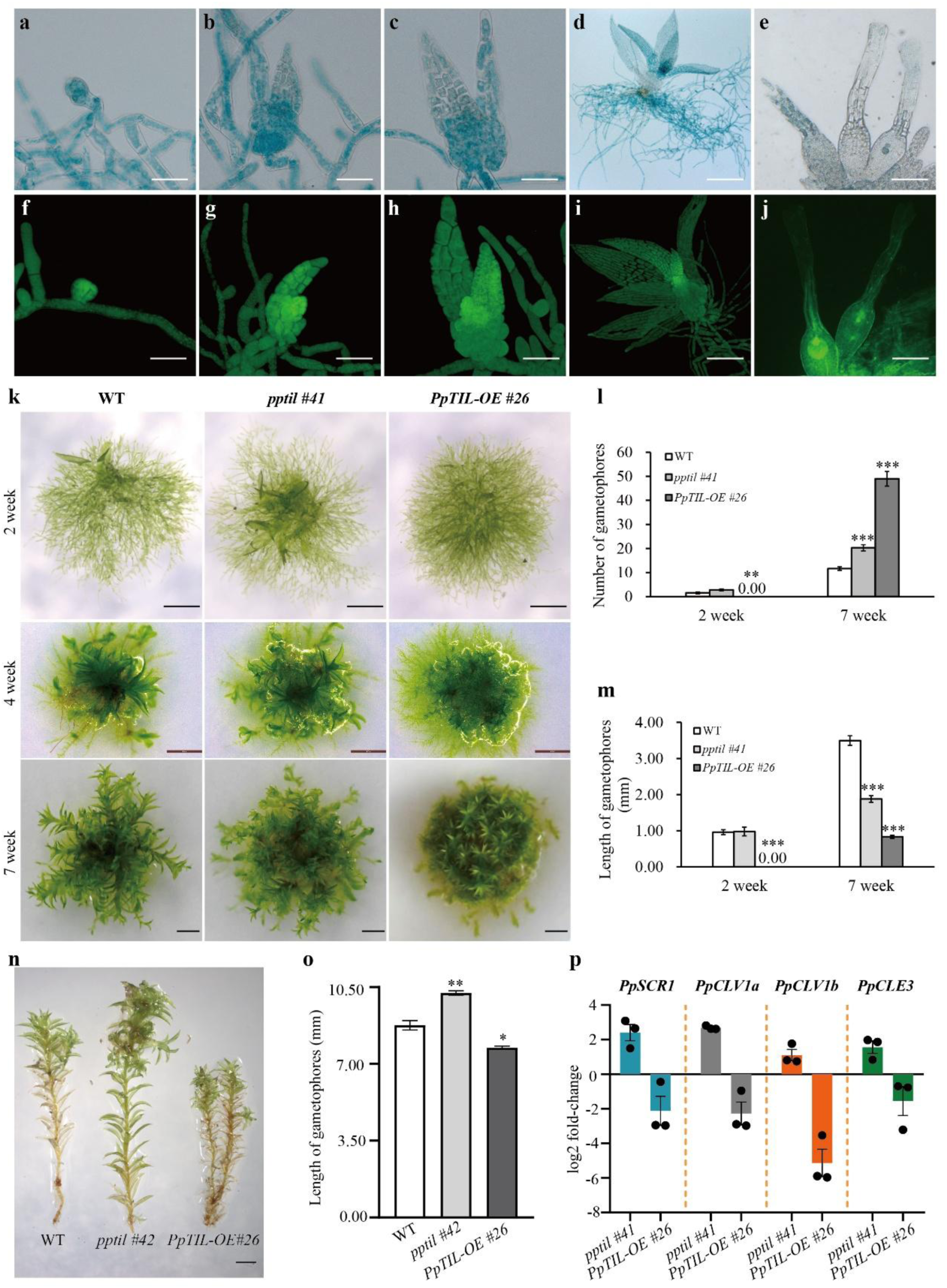
*PpTIL* affects gametophore development in *P. patens*. **a-j,** Expression of *PpTIL* in three-dimensional buds and gametophores based on GUS and GFP signals. **a, f:** buds; **b**-**c, g-h:** juvenile gametophores; **d, i:** leafy gametophores; **e, j:** archegonia. Scale bar: 200 μm**. k,** Gametophore development from 2-to 7-week-old WT, *ko,* and *OE* plants. Protonemata of approximately 1 mm in diameter were transplanted onto BCD medium from 7-day-old plants, and cultivated for two, four, and seven weeks, respectively. Micrograph images provided were observed from ten biological replicates. Scale bar: 1 cm. **l,** Quantification of gametophore number in **k**. **m,** Quantification of gametophore length in **k**. Each bar shows mean ± s.e.m. of at least five biological replicates in **l** and **m**. **n,** Phenotypes of mature gametophores bearing archegonia and antheridia in WT, *ko* and *OE* plants. Scale bar: 1 mm. **o,** Quantification of gametophore length in **n**. Each bar shows mean ± s.e.m. of at least three biological replicates. **p,** Expression profiles of the three-dimensional growth marker gene *PpSCR1*, as well as *PpCLV1a*, *PpCLV1b* and *PpCLE3* that are associated with two-dimensional to three-dimensional transition. Data were collected from three biological replicates. Asterisks indicate a statistically significant difference compared with the wild type based on a two-tailed Student’s *t* test (\**p*< 0.05, \*\**p*< 0.01, *** *p*< 0.001).

To understand how *PpTIL* affects gametophore development, we performed phenotypic analyses of WT, *ko* and *OE* plants under normal growth conditions. After two weeks of cultivation, gametophores became visible in both the wild type and *ko* mutants, with the latter producing more gametophores. Meanwhile, *OE* plants had no gametophore but only dense chloronemata in the middle of colony (Fig. 4k, l). After four weeks, colonies of *OE* lines still had dense chloronemata but gametophores, smaller than those of WT and *ko* plants, emerged on the periphery (Fig. 4k). At seven weeks, both *ko* and *OE* lines had significantly more, but smaller, gametophores than the wild type. Specifically, the number of gametophores increased by about 74% and 320% in *ko* and *OE* plants, respectively, relative to the wild type (Fig. 4k, l), but the length of their gametophores was about 46% and 76% shorter accordingly (Fig. 4k, m). At maturity, gametophores of *OE* lines not only were visibly smaller compared to the wild type, but also senesced early, as evidenced by the many brownish leaves in the lower and middle portion of gametophores (Fig. 4n, o; Supplementary Fig. 8). Conversely, gametophores of *ko* mutants exhibited delayed senescence and were significantly larger at maturity (Fig. 4n, o; Supplementary Fig. 8). Consistent with these observations, chlorophyll (a, b, a+b) and carotenoid contents increased in *ko* mutants but decreased in *OE* lines compared to WT plants (Supplementary Fig. 8).

The above observations (Fig. 4a-d, f-i, k-o) suggest that *PpTIL* affects the transition from two-dimensional to three-dimensional growth and the development of gametophores in *P. patens*. To corroborate these findings, we generated transcriptome data using one-week-old protonemata of WT and transgenic plants, and then examined the expression profile of marker genes for three-dimensional growth of *P. patens*. We found that the expression of SCARECROW transcription factor 1 gene (*PpSCR1*), a known marker of three-dimensional growth in *P. paten*^41^, was significantly increased (over four folds) in *ko* mutants whereas decreased (80%) in *OE* lines compared to WT plants (Fig. 4p). Similarly, *PpCLAVATA1a* (*PpCLV1a*), *PpCLAVATA1b* (*PpCLV1b*) and *PpCLAVATA3-like 3* (*PpCLE3*) in the CLAVATA signaling pathway^40,42,43^, were up-regulated (ca. 2-6 folds) in *ko* mutants but down-regulated (70-97%) in *OE* plants (Fig. 4p). The expression of other known genes related to gametophore development in *P. patens*, such as *DEK1*, *NOG1-2* and *APB1-4*^41,44–46^, changed slightly, but no significant and consistent regulatory pattern was observed between *ko* and *OE* lines. Collectively, these RNA-seq data are consistent with the observation of more gametophores in two-week-old colonies of *ko* mutants, confirming the role of *PpTIL* in gametophore development of *P. patens*.

### *PpTIL* regulates the transition from chloronema to caulonema

Because the initiation of three-dimensional buds occurs in caulonemal cells, the transition from chloronemata to caulonemata represents a key process in gametophore development^12,40^. Therefore, we further compared the protonemal development in WT, *ko* and *OE* plants. Compared to the wild type, an over 45% increase of caulonemal filaments was observed in eight-day-old colonies of *ko* mutants. Conversely, an approximately 60% decrease of caulonemal filaments was found in *OE* lines at the same developmental stage (Fig. 5a, b), in line with the fact that the development of gametophore was delayed in *OE* lines during early stages (Fig. 4k, l). As plants grew, the development of caulonemata became significantly faster in *OE* lines but slower in *ko* mutants. At about twenty days, we observed about the same number of caulonemata in *ko* and WT plants but twice as many caulonemata in *OE* lines (Fig. 5a, c). Compared to WT and *ko* plants, each filament in *OE* lines also developed more caulonemal cells (Fig. 5d). This significantly higher number of caulonemata and caulonemal cells eventually led to more but smaller gametophores at later developmental stages in *OE* lines (Fig. 4k-m).

**Fig. 5.**
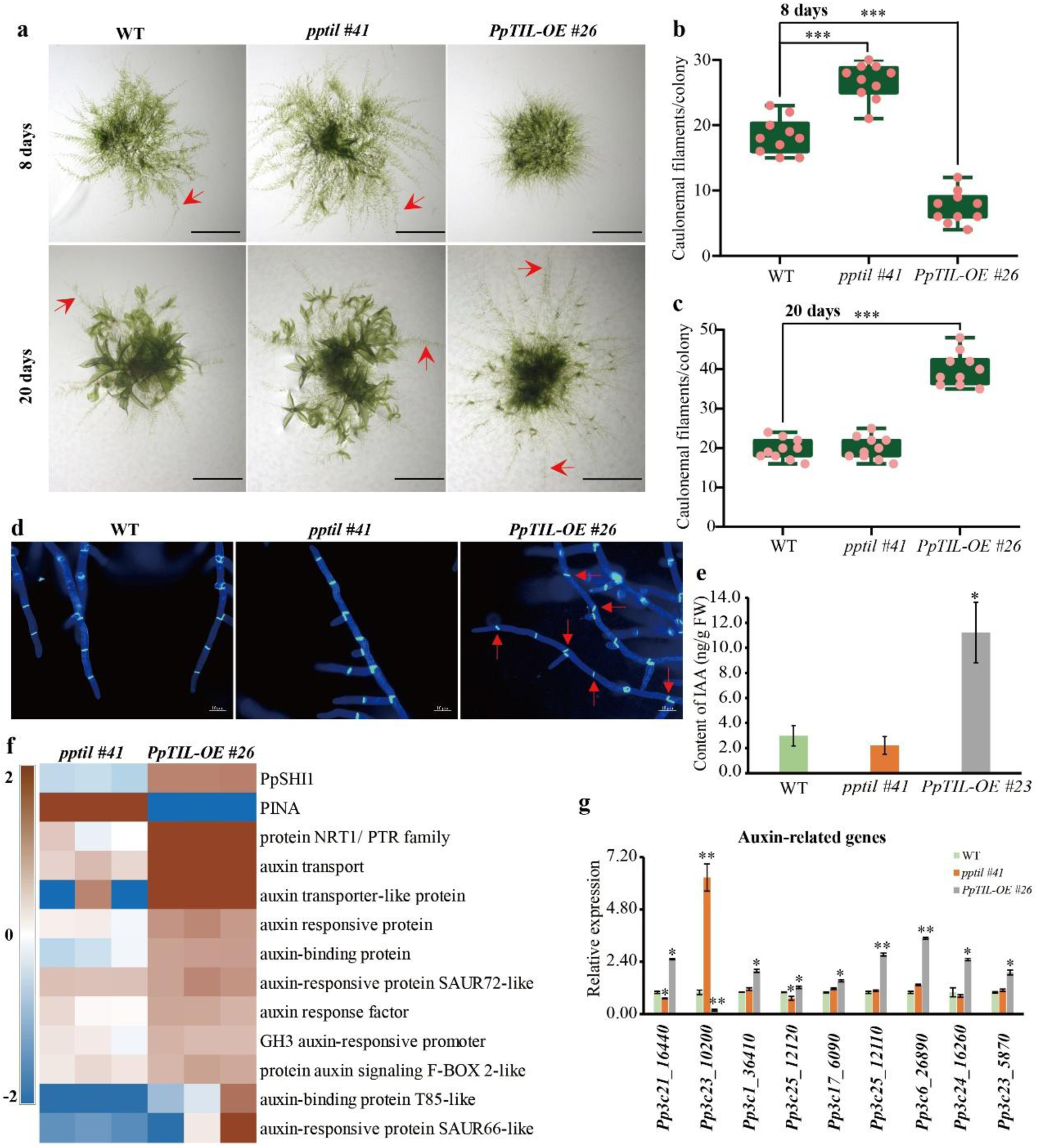
*PpTIL* affects the transition from chloronema to caulonema in *P. patens*. **a,** Development of chloronemata to caulonemata in 8-day-old and 20-day-old WT, *ko*, and *OE* plants, respectively. Protonemata of approximately 1 mm in diameter were transplanted onto BCD medium from 7-day-old plants, and cultivated for eight and twenty days, respectively. Micrograph images provided were observed from ten biological replicates. Scale bar: 500 μm. Red arrowheads indicate caulonemal filaments. **b,** Quantification of caulonemal filaments per colony from 8-day-old WT, *ko*, and *OE* plants. **c,** Quantification of caulonemal filaments per colony from 20-day-old WT, *ko*, and *OE* plants. Each bar shows mean ± s.e.m. of ten biological replicates in **b** and **c**. **d,** Micrographs of calcofluor white (CFW) stained protonemata from 20-day-old WT, *ko,* and *OE* plants. Red arrowheads indicate caulonemal cells. **e,** IAA contents from 10-day-old protonemata of WT, *ko*, and *OE* plants, respectively. **f,** Transcriptional profile from RNA-seq data for a subset of genes related to auxin transport, binding and response in WT, *ko* and *OE* plants. **g,** Expression levels of differentially expressed genes (DEGs) from **f** were confirmed through qRT-PCR. Data are normalized to *PpEF1a*. Each bar shows mean ± s.e.m. of three independent biological replicates. Asterisks indicate a statistically significant difference compared with the wild type based on a two-tailed Student’s *t* test (* *p*<0.05, ** *p*<0.01, *** *p*<0.001). Additional information for each gene is provided in Supplementary Table 1.

The transition from chloronemata to caulonemata is induced by auxin in *P. patens*^47–50^. As such, we measured the auxin content of the WT, *ko* and *OE* plants from 10-day-old protonemata, respectively. We detected no significant difference between *ko* and WT plants but an about two-fold increase in *OE* lines in auxin content (Fig. 5e), in line with the significant increase in the number of caulonemal cells (Fig. 5d). To understand how *PpTIL* affects auxin content, we further analyzed our generated RNA-seq data and identified differentially expressed genes (DEGs) in *ko* and *OE* plants. Notably among the identified DEGs were those related to transport and metabolism of lipids, as well as biosynthesis, transport, binding and response of auxin (Fig. 5f; Supplementary Table 1). Particularly, *PpSHI1*, a positive regulator of auxin biosynthesis^48^, is significantly down-regulated in *ko* mutants but up-regulated in *OE* lines. Conversely, *PpPINA* (*Pp3c23_10200*), a key auxin efflux carrier^51,52^, was significantly up-regulated in *ko* mutants but down-regulated in *OE* lines. The expression of *PpLRL* and *PpRSL,* two bHLH transcription factor genes that promote the chloronema to caulonema transition^47,49^, also increased 3-4 folds in *OE* lines (Supplementary Fig. 9). We further performed qRT-PCR experiments to confirm the expression levels of these genes from RNA-seq data (Fig. 5g; Supplementary Fig. 9).

Because flavonoids negatively modulate the transport of auxin^53–55^, we also analyzed the expression profiles of key genes in flavonoid biosynthesis from transcriptomes of *ko* and *OE* plants. We found that multiple genes in phenylpropanoid and flavonoid biosynthesis, including *phenylalanine ammonia lyase* (*PAL*), *cinnamic acid 4-hydroxylase* (*C4H*), *4-coumarate:CoA ligase* (*4CL*), *flavanone 3-dioxygenase 2-like*, *flavonoid 3’-hydroxylase* (*F3’H*) and *flavonoid 3’5’-hydroxylase* (*F3’5’H*), were significantly up-regulated in *OE* lines, whereas slightly down-regulated in *ko* mutants compared to WT plants (Supplementary Fig. 10; Supplementary Table 2). Considering the changes in the content of auxin and the expression of auxin-related genes in *ko* and *OE* lines (Fig. 5e-g), our data support the earlier suggestion that inhibition of auxin transport by flavonoids disrupts auxin homeostasis in *P. patens*^41,52^.

### *PpTIL* promotes activities of development-related transcription factors

In view of the strong expression of *PpTIL* in apical meristems of gametophores and its roles in auxin biosynthesis, transport and response, we reasoned that *PpTIL* might be involved in stem cell functions and, therefore, investigated whether *PpTIL* affects cell regeneration. Protoplast regeneration was carried out for WT, *ko* and *OE* plants, respectively, and their developmental changes were observed (Fig. 6a). Compared to the wild type, *ko* mutants showed hastened differentiation during the first ten days and, after 10-15 days, bud initiation was observed (Fig. 6b); *OE* lines, on the contrary, exhibited delayed growth during the first ten days of cell differentiation, and then their growth became significantly faster, generating numerous branches and longer cells (Fig. 6a, c), consistent with their dense protonemata during early developmental stages (Fig. 4k).

**Fig. 6.**
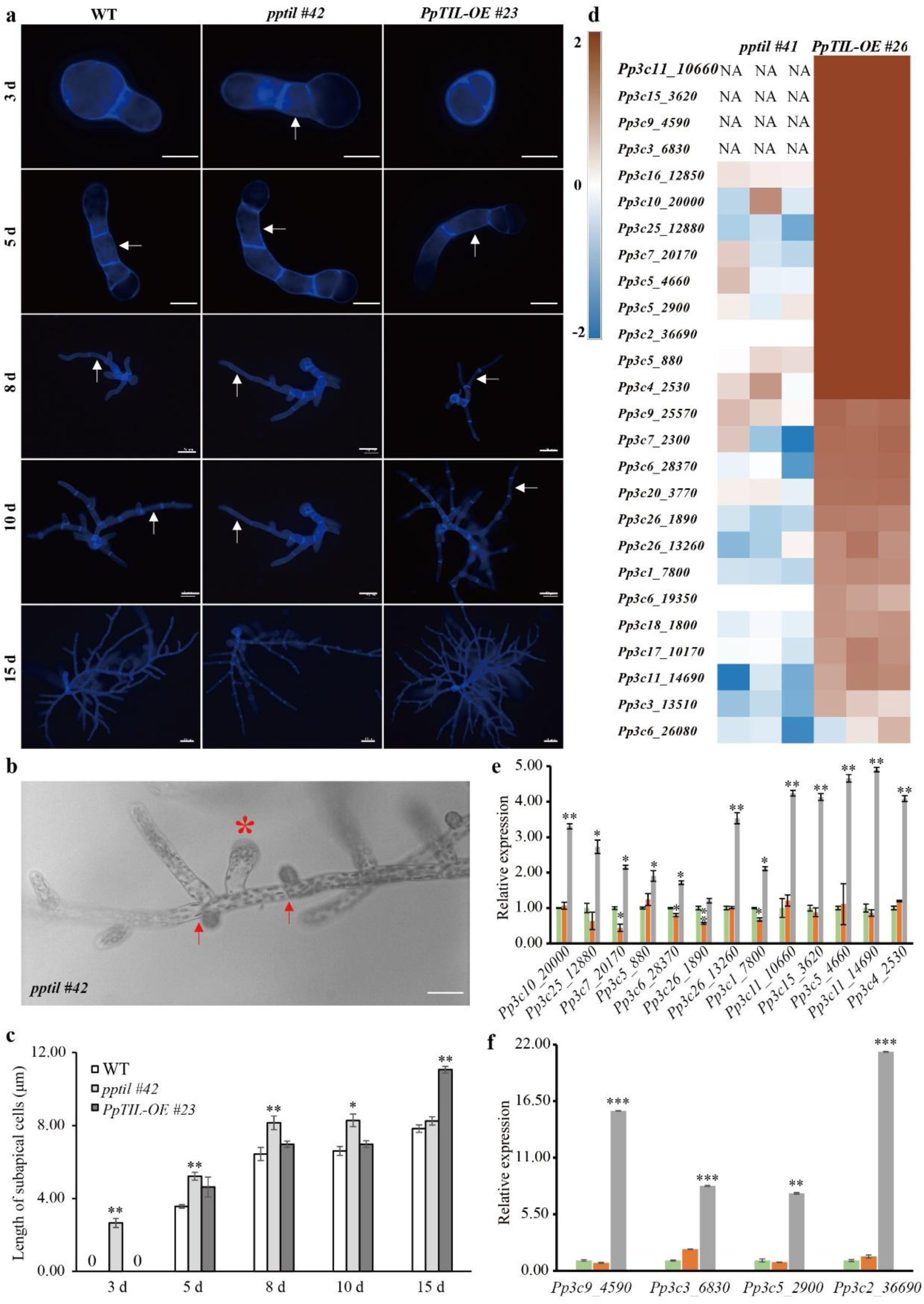
*PpTIL* promotes functions of stem cells and development-related transcription factors. **a,** Protoplast regeneration for WT, *ko*, and *OE* plants, respectively. Microscopic images show protoplast regeneration stained with calcofluor white at 3-15 days, with white arrowheads indicating emerged subapical cells. **b,** Buds in *ko* mutants. Red arrowheads indicate caulonemal cells, and the red asterisk shows an emerged bud. Micrograph image shown was observed from five biological replicates. Scale bars: 10 μm in **a** and **b**. **c,** Lengths of subapical cells from WT, *ko* and *OE* plants shown in **a**. Each bar shows mean ± s.e.m. of three replicates. **d,** Transcriptional profile from RNA-seq data for a subset of genes related to development-related transcription factors in WT, *ko*, and *OE* plants. **e-f,** Expression levels of differentially expressed genes from **d** were confirmed through qRT-PCR. Data are normalized to *PpEF1a*. Each bar shows mean ± s.e.m. of three independent biological replicates. Asterisks indicate a statistically significant difference compared with the wild type based on a two-tailed Student’s *t* test (* *p*<0.05, ** *p*<0.01, *** *p*<0.001) in **c**, **e** and **f**. Additional information for each gene is provided in Supplementary Table 3.

The above observations not only support the role of PpTIL in stem cell functions, but also are consistent with our generated RNA-seq data, which included various development-related transcription factors that were differentially expressed in *ko* and *OE* plants. Notably, APETALA2 (AP2) and TINY-like were down-regulated in *ko* mutants and up-regulated in *OE* lines (Fig. 6d; Supplementary Table 3). AP2 transcription factors are known to regulate stem cell functions, cell reprogramming and gametophore formation in *P. patens*^46,56^. In *Arabidopsis*, TINY affects plant growth and development by perceiving brassinosteroid (BR) signal, and *TINY-OE* plants exhibit stunted growth^57^. The expression of the above transcription factor genes was further confirmed by qRT-PCR experiments (Fig. 6e, f). Overall, the above findings suggest that PpTIL enhances functions of development-related transcription factors and activities of stem cells.

### *PpTIL* interacts with proteins for lipid metabolism and other activities

Lipocalins are known to participate in lipid transport and metabolism, as well as miscellaneous other activities^15,58^. As expected, our RNA-seq data showed that certain lipid metabolism-related genes were differentially expressed in *ko* and *OE* plants (Fig. 7a; Supplementary Table 4). Additional enrichment analyses of DEGs identified pathways related to metabolism of lipids and their derivatives, particularly those related to ether lipids, glycerolipids, glycerophospholipids, alpha-linolenic acids, cutin, suberine and wax (Fig. 7b; Supplementary Table 5). Intriguingly, several key genes in the biosynthesis of lipid-derived oxylipin 12-oxophytodienoicacid (OPDA), including linoleate 9S-lipoxygenase (*LOX*, *Pp3c17_16260*), allene oxide synthase (*AOS*, *Pp3c7_25640*) and allene oxide cyclase (*PpAOC1*: *Pp3c4_22490*, *PpAOC2*: *Pp3c5_3730*, and *Pp3c2_24500*), were significantly up-regulated in *OE* lines (Supplementary Fig. 11). The above finding is consistent with the observation that PpTIL is localized in chloroplasts (Fig. 2), where fatty acids are produced and a major site of lipid biosynthesis^59,60^.

**Fig. 7.**
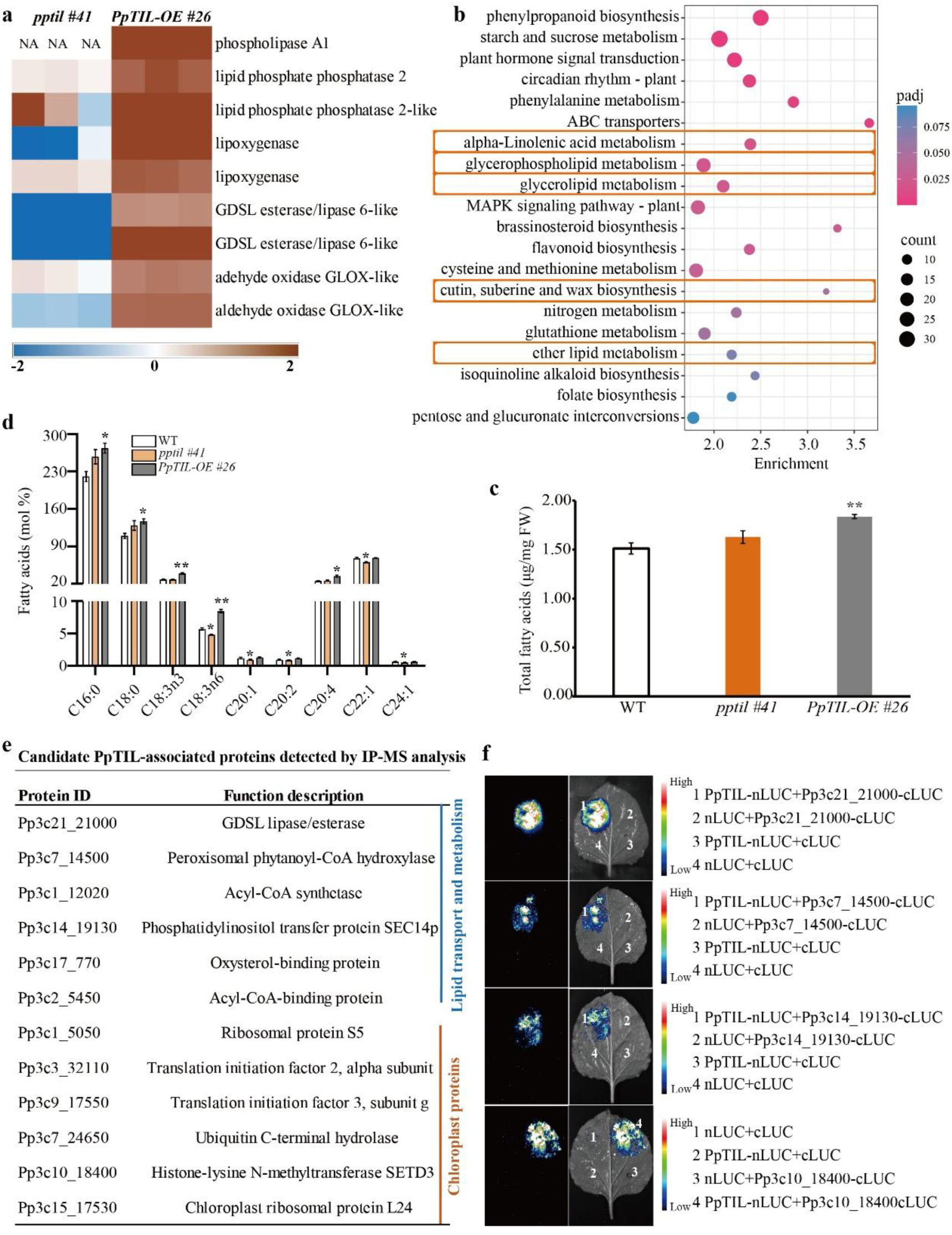
*PpTIL* is involved in the transport and metabolism of lipids and other activities in *P. patens.* **a,** Transcriptional profile from RNA-seq data for differentially expressed genes (DEGs) related to lipid metabolism in *ko* and *OE* plants compared to the wild type. Additional information for each gene is provided in Supplementary Table 4. **b,** Kyoto Encyclopedia of Genes and Genomes (KEGG) analysis of the identified DEGs. Orange frames are related to lipid metabolism. Additional information is provided in Supplementary Table 5. **c,** Total fatty acids content from 10-day-old protonemata of WT, *ko,* and *OE* plants, respectively. **d,** Contents of saturated and unsaturated fatty acids from WT, *ko* and *OE* plants. Each bar shows mean ± s.e.m. of three independent biological replicates. Asterisks indicate a statistically significant difference compared with the wild type based on a two-tailed Student’s *t* test (* *p*<0.05, ** *p*< 0.01). **e,** PpTIL-associated candidate proteins detected by IP-MS analysis using GFP-tag of *PpTIL*pro*:PpTIL-EGFP-GUS* plants. **f,** Luciferase complementation imaging (LCI) analysis of interactions between PpTIL and candidate proteins through transient expression in tobacco leaves. The N-terminus of LUC (nLUC) was fused with PpTIL, and C-terminus of LUC (cLUC) was fused with a candidate protein. The presence of signal indicates that two proteins attached to the N- and C-termini can interact.

We further analyzed the changes in fatty acid content between *ko* or *OE* and WT plants. Compared to the wild type, the total fatty acids were over 20% higher in *OE* plants but not significantly different in *ko* mutants (Fig. 7c). Importantly, palmitic acid (16:0), stearic acid (18:0), and linolenic acid (18:3n3 and 18:3n6), involved in membrane lipid remodeling such as changes in fatty acid composition and fatty acid desaturation^60,61^, were significantly increased in *OE* lines (Fig. 7d). Moreover, very long-chain polyunsaturated fatty acids (e.g., C20:1, C20:2, C22:1 and C24:1), which are essential for bryophytes to survive and adapt to harsh environment^62^, were significantly decreased in *ko* mutants (Fig. 7d). These results are consistent with the observations that *PpTIL* promotes tolerance to abiotic stresses in *P. patens* (Fig. 3). Because lipid metabolism is closely tied to metabolism of carbohydrates and flavonoids^63–65^, we also evaluated the expression of genes related to the above metabolic pathways. Compared with the wild type, many of these genes was up-regulated by at least two folds in *OE* lines, while slightly down-regulated in *ko* mutants (Supplementary Figs. 10, 12). Consistent with the above data, differentially expressed metabolites from *ko* and *OE* plants were enriched in flavonoid biosynthesis, carbohydrate metabolism, and related pathways (Supplementary Fig. 13). Altogether, our results confirmed the role of PpTIL in fatty acid and lipid metabolism, as well as their effects on related processes.

We performed Immunoprecipitation-Mass Spectrometry (IP-MS) assays using GFP tagged transgenic plants to search for possible proteins interacting with PpTIL. Candidate interacting proteins of PpTIL identified from the IP-MS data were mostly related to lipid transport and metabolism, as well as chloroplast activities (Fig. 7e). We further performed split-luciferase complementary assays to validate a subset of candidate proteins. As a result, we found that PpTIL interacts with GDSL lipase/esterase (Pp3c21_21000), phytanoyl-CoA hydroxylase (PAHX, Pp3c7_14500), SEC14-like phosphatidylinositol transfer protein (SEC14L-PITP, Pp3c14_19130), and Rubisco methyltransferase (Pp3c10_18400), respectively (Fig. 7f). Among these interacting proteins, GDSL lipases and esterases hydrolyze a wide variety of lipids and other substrates^66,67^, whereas PITPs can transport and exchange various phospholipid species (e.g., phosphatidylinositol, phosphoinositides and phosphatidic acids) and other lipophilic molecules between membranes or cellular compartments^68–70^. The other two interacting proteins are involved in activities related to chloroplasts. Specifically, Rubisco methyltransferase is localized in chloroplast stroma in *Arabidopsis*, where it protects grana by suppressing stress response inducted by singlet oxygen^71^. PAHX is involved in the metabolism of phytol generated from chlorophyll degradation^72^. Overall, these data provide additional evidence that *PpTIL* is involved in other cellular activities in *P. patens*, in addition to lipid transport and metabolism.

## Discussion

Despite extensive research over the years, many of the land plant innovations and adaptive strategies remain to be fully understood. Lipocalins are considered to be “optimizers” for organismal development and adaptation^58^, and they are known to play a critical role in membrane maintenance, metabolic regulation, antioxidation, sexual reproduction, and many other physiological and developmental processes in animals and bacteria^15,17,23,58^. Plant lipocalins are often more similar to bacterial and certain animal sequences, and they were thought to be derived from the cyanobacterial progenitor of plastids^25^. Our data show that at least three lipocalin groups exist in land plants, including TILs, CHLs, and a distinct clade uncovered in this study that is specific to seedless plants (SLLs) (Fig. 1c, d), none of which is specifically related to cyanobacterial sequences. Admittedly, the evolutionary history of lipocalins is very complex, but still both sequence comparisons and phylogenetic analyses point to a bacterial affinity of plant TILs (Fig. 1b, c). We speculate that plant *TIL* genes were horizontally acquired from bacteria by the ancestral streptophyte, similar to many other genes that were transferred from microbes to early land plants and their charophyte relatives^73–76^. The SLL sequences are predicted to be secreted (extracellular) proteins (Fig. 1a) and have apparently been lost in seed plants. Although the exact functions and involved processes of SLLs remain unclear, the loss of genes from seed plants, particularly those horizontally acquired or related to stress tolerance, have also been documented in multiple other studies^75,77–79^.

Our data suggest that PpTIL is a versatile regulator for various physiological and developmental processes. Compared with TILs in flowering plants, which are primarily known to be localized at the plasma membrane and involved in abiotic stresses^24,26,80^, PpTIL not only is functionally conserved in stress response (Fig. 3), but also affects protonemal and gametophore development (Figs. 4-5). In addition, given its strong expression in archegonia (Fig. 4e, j; Supplementary Fig. 7), we speculate that PpTIL is also involved in sexual reproduction and related processes. This versatility is partially reflected in the subcellular localization of PpTIL both at the plasma membrane and in chloroplasts. While the plasma membrane protects cells from stress damage and regulates external signal perception and intercellular exchange^81,82^, chloroplast activities are intertwined with numerous other cellular activities in addition to carbon fixation. Consistent with these findings, both protein-protein interaction and lipodomic data indicate the functional association of PpTIL with chloroplast activities and lipid metabolism. Particularly, protein-protein interaction data show that PpTIL binds not only with some key players in lipid signaling and metabolism (e.g., SEC14L-PIPT and GDSL lipase/esterase)^69,70^, but also with homologous proteins that affect chloroplast grana protection (i.e., Rubisco methyltransferase) and chlorophyll catabolism (i.e., PAHX), in line with its versatile role in physiological and developmental processes.

The versatile role of PpTIL in various physiological and developmental processes is not entirely surprising, particularly given the extensive similar reports of lipocalins in animals^15,21,23^, where they are known to participate in numerous activities or considered to be moonlighting proteins^23,58,83^. For instance, human ApoD can form different oligomers (monomers, dimers, and tetramers) or associate with different lipid-rich structures, thus modulating a wide variety of cellular activities^23,84^, presumably due to the intrinsic binding property of lipocalin pocket and the ability to protect membranes against oxidative stress^23^. At present, the detailed biochemical mechanisms for the involvement of lipocalins in physiological and developmental processes of land plants remain largely unclear. In flowering plants, available data support a role of lipocalins in binding to lipids and related molecules (e.g., non-specific lipids and retinal)^28,33^, as well as in antioxidative activities^26–29^. In our study, in addition to its role in lipid metabolism and chlorophyll catabolism, PpTIL might also participate in other activities, suggesting a far greater role of lipocalins in plants than currently realized. Notably, PpTIL overexpression significantly up-regulates the expression of biosynthetic genes (e.g., *LOX*, *AOCs*; Supplementary Fig. 11) of the oxylipin OPDA, a precursor of bioactive jasmonate in bryophytes^85^ and a signaling molecule in plant defense and development^86,87^, and PpTIL interacts with PIPT (Fig. 7f), a key player in the transport of phospholipid signaling molecules, suggesting that PpTIL is likely involved in lipid signaling pathways in *P. patens*. Several other activities, from auxin biosynthesis and transport (Fig. 5e-g) to activities of development transcription factors (Fig. 6d-f), were also changed in *PpTIL ko* and *OE* lines. Some of these changes are consistent with earlier reports, and they are likely affected indirectly by PpTIL. For instance, the negative correlation between auxin content and flavonoids in *PpTIL ko* and *OE* plants, as revealed in this study, is likely caused in part by the inhibitory effects of flavonoids on auxin polar transport in plants^53,55,88^. Nonetheless, plant lipocalins are vastly understudied, particularly in bryophytes, and detailed investigations are needed to fully understand their role in the physiology and development of land plants.

The finding that PpTIL is involved in various physiological and developmental processes also raises a question about the role of lipocalins in plant colonization of land. Major physiological and developmental innovations to overcome challenges in terrestrial environments, such as stress responses and three-dimensional growth^4,6^, were key to the establishment of plants on land. These innovations allowed early plants to tolerate heat, cold, drought and related stresses, but also to thrive and diversify in various terrestrial settings. The involvement of PpTIL in abiotic stress response and three-dimensional development in *P. patens* (Figs. 3-6) suggests a pivotal role of lipocalins in early land plant evolution. Notably, both TILs and SLLs are largely restricted to land plants and their charophyte algal relatives. Particularly, SLLs are retained only in charophytes and seedless plants (Fig. 1c, d). Despite the lack of functional information on SLLs, this unique retention pattern in early diverging land plants and their close relatives points to the importance of SLLs during the transition of green plants from water to land. Further investigations on these *SLL* genes in bryophytes and charophytes should provide a better understanding of the role of lipocalins in plant colonization of land.

## Materials and Methods

### Plant material and growth conditions

The *Physcomitrium patens* ‘Gransden 2004’ was used as WT plants in this study. Protonemata were grown on BCDAT medium, and gametophores were grown on BCD medium (minus ammonium tartrate), at 25°C under a 16-hour light and 8-hour dark regime, and a light intensity of 80 µmol photons m^−2^ s^−1^.

### Identification of lipocalins and phylogenetic analyses

TIL (AT5G58070, Pp3c18_22450) and CHL (AT3G47860, Pp3c9_6110) protein sequences from *A. thaliana* and *P. patens* were used as queries to perform BLAST searches against NCBI non-redundant (nr) protein sequences, OneKP, published plant genomes that are not included the nr database, over 160 unpublished bryophyte genomes generated by our collaborators, and other related sequence databases (E-value cutoff=1e-4). To identify remote homologs, additional HMMER searches were performed with lipocalin and lipocalin-like domains (PF00061 and PF08212) as query. Sequences were sampled from representative groups of all three domains of life (bacteria, archaea, and eukaryotes) and viruses for phylogenetic analyses. Multiple sequence alignments were carried out using Mafft v7.490, with default settings, followed by manual removal or trimming of gaps and ambiguously aligned sites. Only the lipocalin and lipocalin-like domain regions were used for phylogenetic analyses. Phylogenetic analyses were performed using the maximum likelihood method implemented in IQ-Tree v2.1.4, with the best-fitting amino acid substitution model (Q.pfam +G4) selected based on the Bayesian Information Criterion (BIC). Branch support values were estimated with 1,000 replicates using ultrafast bootstrapping in IQ-Tree.

### Measurement of gametophores and caulonemal filaments

Seven-day-old protonemata of approximately 1 mm in diameter were transplanted onto BCD medium and cultivated under normal growth conditions. The number of gametophores was determined by counting all gametophores per colony after 2-7 weeks. Meanwhile, lengths of gametophores in each colony at different developmental stages were measured. The transition from chloronemata to caulonemata was quantified by counting caulonemal filaments per colony of 8- and 20-day-old plants, respectively.

### Plasmid construction for knockout (*ko*) and overexpression (*OE*)

The vector pTN182 was used to delete *PpTIL* from the wild type. Genome fragments containing upstream (1,084 bp) and downstream (825 bp) flanking regions of *PpTIL* for homologous recombination were inserted into the pTN182 vector via restriction enzymes KpnI-SalI and XbaI-BamHI, respectively (Supplementary Fig. 4). Primers used for constructing the gene knockout vector are provided in Supplementary Table 6.

The vector pPOG1 containing upstream and downstream recombinant fragments *PIG1bL* and *PIG1bR* (Supplementary Fig. 5) was used, respectively, to construct overexpression plasmid. *PpTIL* coding sequence (CDS) without the stop codon was amplified from cDNA by RT-PCR using high-fidelity enzymes, and then cloned into the pPOG1 vector through NotI and XhoI restriction enzymes. Primers used for *PpTIL* CDS amplification are listed in Supplementary Table 6.

### Protoplast transformation

To obtain *PpTIL ko* and *OE* lines in *P. patens*, polyethylene glycol (PEG)-mediated protoplast transformation was performed^37^. Protoplasts were first prepared using 1-2% driselase, and the final concentration of 1.6 × 10^6^ protoplasts per ml of solution was obtained. Approximately 30 μg of DNA fragment was used to incubate 300 μl of the prepared protoplasts under PEG induction condition, followed by a heat shock at 45°C for 5 minutes, then recovered in water bath at room temperature for 10 minutes. Subsequently, the transformed protoplasts were equilibrated by 8% mannitol, and cultured on bottom medium (BCDAT medium supplemented with 6% mannitol and 10 mM CaCl2) with cellophane for about a week. All transgenic lines were selected by nonselective media and selective media containing 20 μg ml^−1^ G418 or 20 μg ml^−1^ hygromycin, and then screened using genomic PCR and real-time quantitative RT-PCR (qRT-PCR). Additionally, flow cytometry was used to analyze the ploidy of all transgenic plants.

### Chromosome ploidy analyses

The chromosome ploidy of *ko* and *OE* plants was analyzed, respectively, using flow cytometer (BD FacsCalibur). Protonemata were chopped and incubated with propidium iodide (PI), and the relative fluorescence intensity of stained nuclei was measured. The flow cytometer used an argon ion laser with an excitation wavelength of 488 nm. The emission spectrum of PI was detected at 620 nm by fluorescence-2 (FL2) filter. The flow rate was set at around 100 nuclei per second, and fluorescence was collected from at least 6,000 cells for each sample. Parameters were kept constant for all samples. BD CellQuest was used to acquire and analyze the generated data. Histograms were analyzed using ModFit 3.0.

### Protein subcellular localization

The vector pTN85 was used to generate stable transgenic plants with green fluorescent protein (GFP) and β-glucuronidase (GUS) tags. The upstream (1,033 bp) and downstream (905 bp) flanking regions of *PpTIL* were amplified using the genomic DNA of *P. patens* as template, and then inserted into the pTN85 vector via restriction enzymes KpnI-XhoI and BamHI-XbaI, respectively (Supplementary Fig. 3). *PpTIL*pro*:PpTIL-EGFP-GUS* stable lines were obtained through PEG-mediated protoplast transformation. Primers used for vector construction are provided in Supplementary Table 6.

*PpTIL*pro*:PpTIL-EGFP-GUS* plants were grown on BCD and BCADT media, respectively. GFP signals were observed for protonemata and gametophores at different developmental stages. The GFP fluorescence was excited at 488 nm, and images were captured using a microscope (Zeiss LSM 900, Germany).

### Histochemical GUS assays

*PpTIL*pro*:PpTIL-EGFP-GUS* plants containing GUS tag were cultured on BCD medium, and incubated in 20 μl 1× GUS solution for reaction at 37°C. The reaction time varied depending on the expression level of GUS in the detected tissues. Histochemical GUS assays were performed using a GUS stain kit (Real-Times, China), and the preparation of the 1× GUS solution followed the manufacturer’s instructions. In addition, plant samples were treated with absolute ethyl alcohol to remove pigments. The stained tissues were observed and photographed using a light microscope (Zeiss Scope.A1, Germany).

### Total RNA extraction and qRT-PCR analyses

Approximately one-week-old protonemata were collected and ground in liquid nitrogen. About 100 mg samples were used for total RNA extraction using Omni plant RNA kit (CWBIO) and DNase I (Solarbio), following the provided extraction protocol. RNA was reverse transcribed into cDNA by M-MLV reverse transcriptase (Accurate Biology) using oligo dT primers and dNTP substrates.

The synthesized cDNA was used as the template of qRT-PCR. The reaction mix was prepared following the protocol of SYBRPrime qPCR kit (Fast HS) (Biogrund), and qRT–PCR was performed using Bio-Rad CFX96 Real-Time System (Thermo Fisher Scientific), with three independent biological replicates. The melting curve was added for primers used for the first time to facilitate specificity analyses. The relative expression of genes was calculated based on the number of cycles (Cq value). *PpEF1α* (elongation factor 1-alpha, *Phypa_439314*) was used to normalize the data in *P. patens*.

### Transcriptome sequencing and analyses

Total RNA from one-week-old protonemata of *P. patens* was extracted using the Omini plant RNA kit (CWBIO) and DNase I (Solarbio). Library construction was then performed using the NEBNext® Ultra™ RNA Library Prep Kit for Illumina®, and sequencing was conducted using Illumina NovaSeq 6000 (Illumina, USA). The obtained reads were filtered and then aligned with the reference genome of *P. patens* from Phytozome (https://phytozome.jgi.doe.gov) using Hisat2. Differentially expressed genes (DEGs) in *ko* and *OE* plants compared to the wild type were identified using DESeq2^89^. Enrichment analyses of Gene Ontology (GO) and Kyoto Encyclopedia of Genes and Genomes (KEGG) pathways for DEGs were performed using ClusterProfiler.

### Measurement of IAA levels

Ten-day-old protonemata of WT, *ko*, and *OE* plants were collected to measure the content of indole-3-acetic acid (IAA), using ultra-high performance liquid chromatography (Vanquish, UPLC, Thermo, USA) and high-resolution mass spectrometry (Q Exactive, Thermo, USA). The mass spectrometry data were processed using TraceFinder, and quantitative analysis was performed using the external standard method, with three replicates for each sample.

### Analyses of fatty acids content

Protonemal tissues were collected from WT, *ko*, and *OE* plants that had been cultured for ten days on BCDAT medium. Gas chromatography-tandem mass spectrometry (Agilent Technologies Inc. CA, UAS) was used to detect fatty acids content. Quantitative analyses were performed using the external standard method, with three independent biological replicates.

### Lipidomic analyses

Protonemata from 10-day-old WT, *ko* and *OE* plants were used for lipidomic analyses. Lipids were extracted using chloroform/dichloromethane/methanol (2:1:1), and then detected using ultra-high performance liquid chromatography (Vanquish, UPLC, Thermo, USA) and high-resolution mass spectrometry (Q Exactive HFX, Thermo, USA), with three independent biological replicates for each sample. The generated data were analyzed using Quant-My-Way (Agilent Technologies), and the external standard method was used for quantitative analyses. Differentially expressed metabolites (DEMs) in *ko* and *OE* plants were identified using partial least squares discriminant analysis (PLS-DA) and orthogonal partial least squares discriminant analysis (OPLS-DA), followed by GO and KEGG pathway enrichment analyses of identified DEMs.

### Immunoprecipitation-Mass Spectrometry (IP-MS) analyses

Protonemata of *PpTIL*pro*:PpTIL-EGFP-GUS* and WT plants were ground in liquid nitrogen, and total protein was extracted using immunoprecipitation (IP) buffer (50 mM Tris-HCl, pH 7.4, 150 mM NaCl, 1 mM EDTA-2Na, 5% glycerol, 5% NP-40 and 1× protease inhibitor cocktail). Extracted proteins were incubated with GFP-Trap magnetic beads (Epizyme) for 2-4 hours at 4°C. After incubation, the beads were washed three times with washing buffer (50 mM Tris-HCl, pH 7.4, 150 mM NaCl, 1 mM EDTA-2Na, 5% glycerol and 0.05% Tween 20). Proteins on beads were eluted by boiling and analyzed using mass spectrometry.

### Luciferase complementation imaging assays

The cDNA of *PpTIL* was cloned into pCAMBIA1300-nLUC (Takara, Japan) by restriction enzymes KpnI and SalI. The full-length cDNAs of *Pp3c21_21000*, *Pp3c7_14500*, *Pp3c14_19130* and *Pp3c10_18400* were cloned into pCAMBIA1300-cLUC (Takara, Japan), respectively, via restriction enzymes KpnI and SalI. All constructed plasmids were transformed into *Agrobacterium tumefaciens* EHA105, and an equal volume mixture of *A. tumefaciens* containing the nLUC and the cLUC fusion plasmids was injected into *Nicotiana benthamiana* leaf epidermal cells. Luciferase signals were detected using an automatic chemiluminescence image analysis system (Tanon 520, China) after incubation with luciferin (100 μM) in the dark. Primers used to amplify cDNA of related genes are provided in Supplementary Table 9.

### Calcofluor white staining

Protoplasts and protonemata were incubated with calcofluor white (CFW) staining solution for 5 minutes using a plant cell wall CFW staining kit (Genmed, China), and then washed in a cleansing solution (Reagent A) with gentle shaking. Images were obtained at 355 nm excitation and 440 nm emission using a light microscope (Zeiss LSM 900, Germany).

### Histochemical staining for ROS detection

Protonemata were immersed in 1 ml of 0.1% (w/v) NBT (Coolaber, SL18061) and 1 mg/ml DAB (Coolaber, SL1805) staining solution, respectively. Samples were under vacuum treatment for 30 minutes, followed by incubation in NBT staining solution for 60 minutes and DAB staining solution overnight at room temperature, respectively. The stained samples were treated with absolute ethyl alcohol to remove pigments, and then photographed using a light microscope (Zeiss Scope.A1, Germany).

### Immunoelectron microscopy analysis

Plant samples of *PpTIL*pro*:PpTIL-EGFP-GUS* were fixed in glutaraldehyde and then embedded in resin. Ultra-thin slices (70 nm thickness) from samples were prepared with an ultra-microtome (Leica EM UC7) and mounted on 100 mesh gold transmission electron microscope grids. The grids were blocked in PBS (1% BSA, 0.05% Triton-X100, 0.05% Tween-20, PH7.4) for 5 minutes at room temperature, followed by incubation with anti-GFP antibody (1:100 dilution) in PBS (1% BSA, 0.05% Tween-20, PH7.4) overnight at 4°C. The grids were then washed six times (5 minutes/wash) with PBS and incubated with 10 nm colloidal gold-conjugated goat anti-rabbit IgG (1:100 dilution, Bioss) at room temperature for 1 hour. Sample grids were then rinsed six times (5 minutes/wash) with PBS and four times (5 minutes/wash) with distilled water, followed by staining with uranylacetate and observation using a transmission electron microscope at 80 kV (Thermo Scientific™ Talos™ L120C).

## Data availability

Data supporting the findings of this study are available from the corresponding author upon request.

## Acknowledgments

We thank Jill Harrison for comments to improve the manuscript, Yanxia Jia for chromosome ploidy analyses, and Zhijia Gu for technical support of microscopy. The work is funded in part by Yunnan Revitalization Talent Support Program “Young Talent” Project to SW.

## Author contributions

SW and JH conceived and designed the study. SW, JM and JH performed experiments and data analyses. QW, YG, XH and HS participated in experiments and data analyses. SW, JM and JH wrote the manuscript.

