## Supplementary Fig for "Lipocalins are versatile regulators of development and stress response in mosses"

**for**

**Wang et al.**

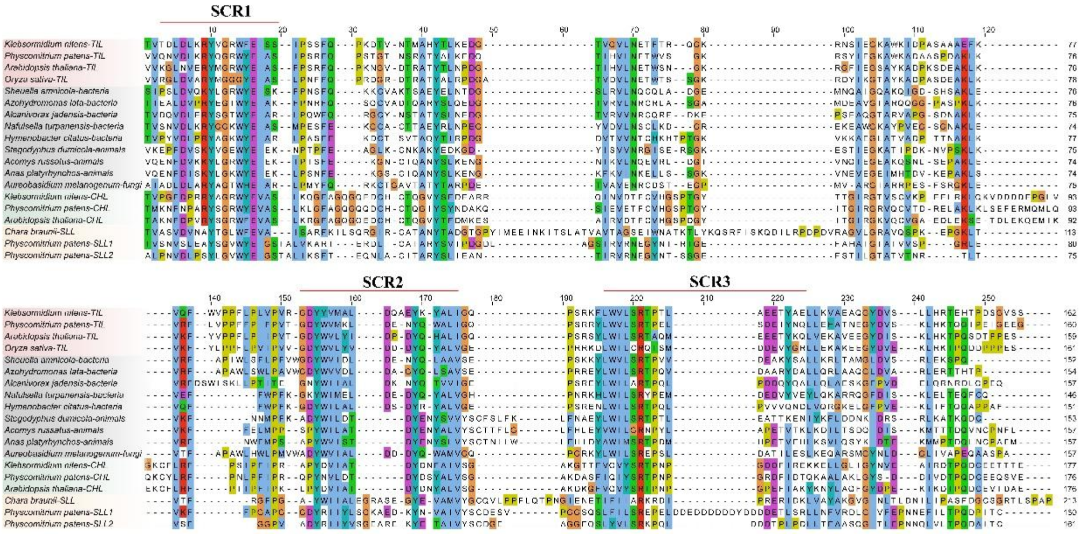


**Supplementary Fig. 1.** Multiple sequence alignment of plant lipocalins and related sequences from other organismal groups. Only conserved regions are displayed. Three main structurally conserved regions (SCRs) are marked.

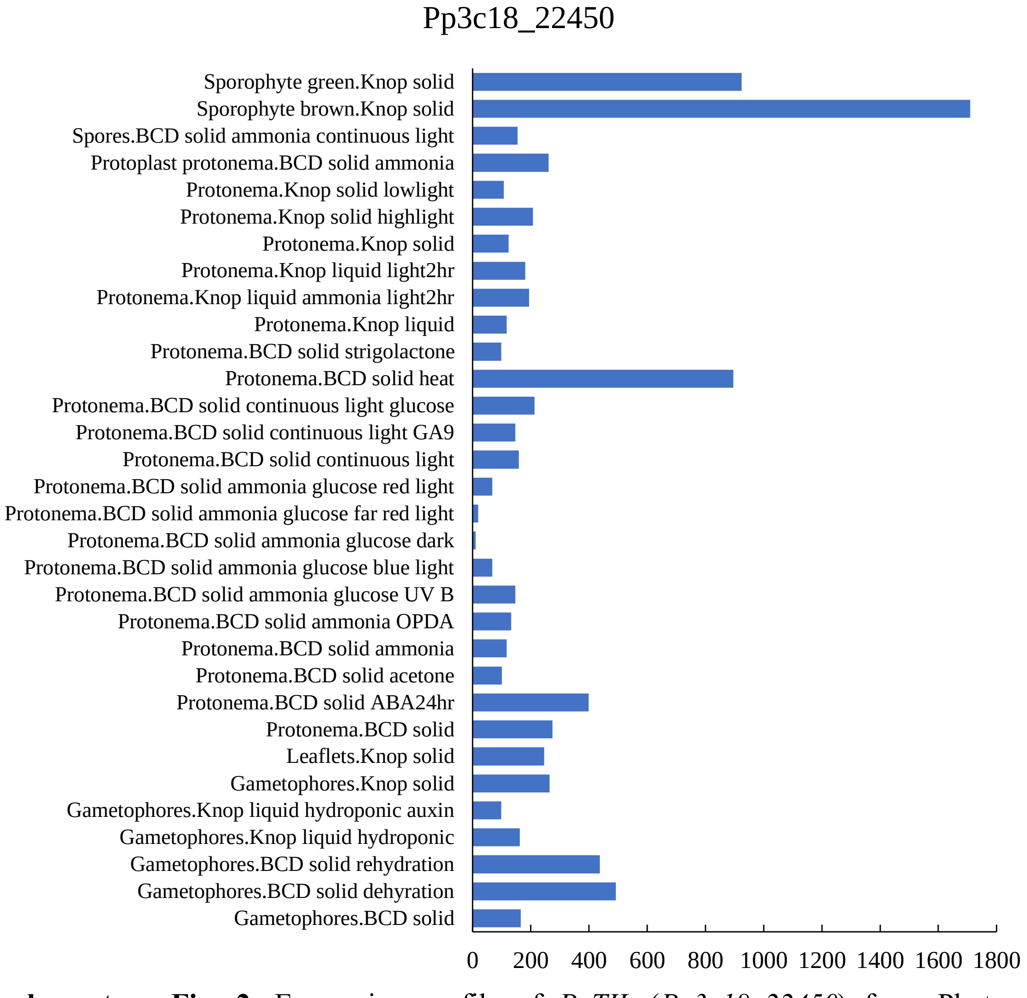


**Supplementary Fig. 2.** Expression profile of *PpTIL* (*Pp3c18_22450*) from Phytozome database.

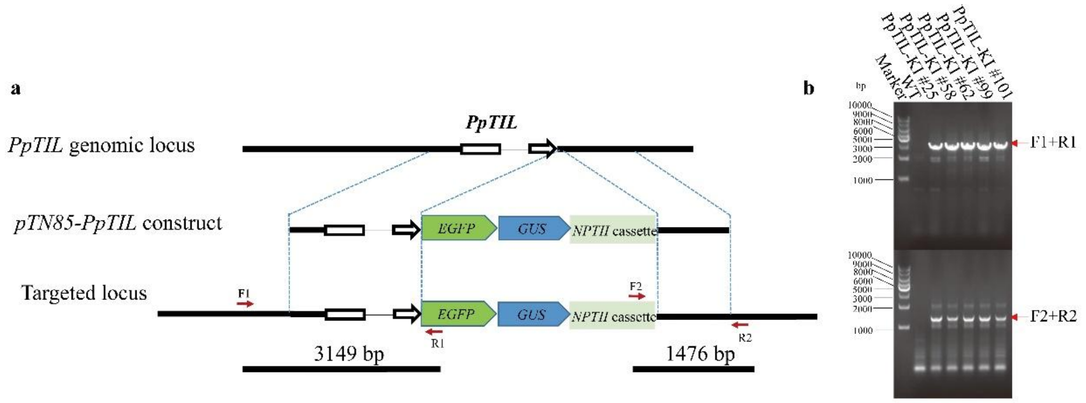


**Supplementary Fig. 3.** Genotyping of *PpTIL* knockin double tag lines. **a,** Construction of *PpTIL* knockin double tag lines. White boxes show the exons of *PpTIL*, and red arrows denote positions of primers used for genomic PCR. **b,** Genomic PCR for *PpTIL* knockin double tag lines. The fragment length of genomic PCR is consistent with expectation as shown in **a**.

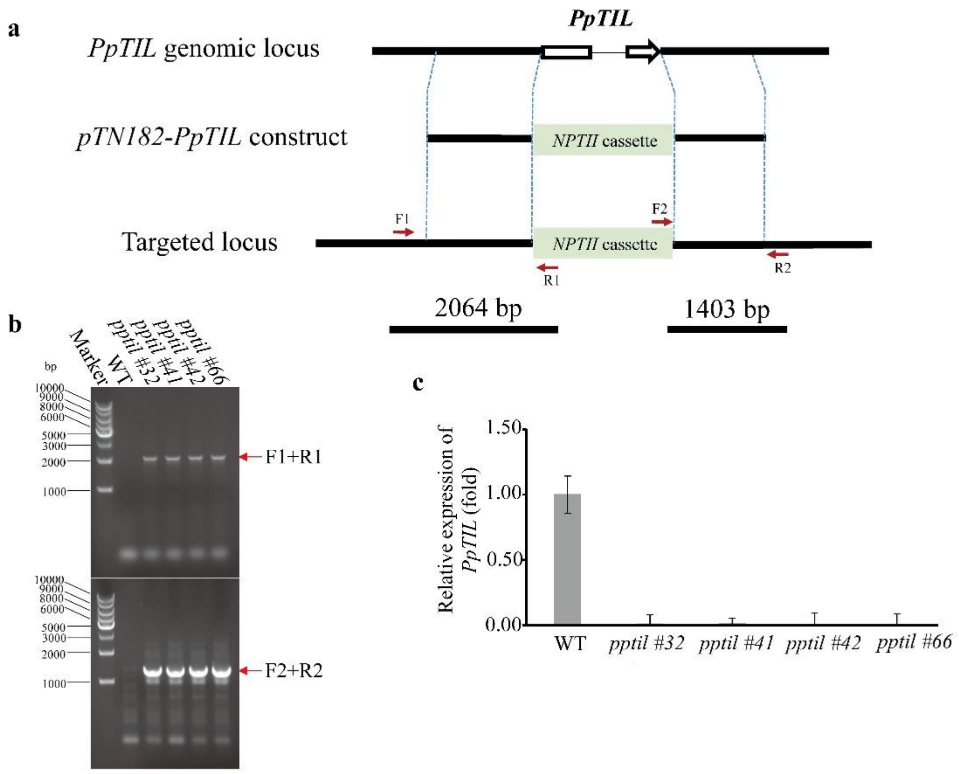


**Supplementary Fig. 4.** Generation and molecular identification of *PpTIL ko* mutants. **a,** Schematic diagram of *PpTIL ko* construction and primers used for genotyping. White boxes indicate exons of *PpTIL*. The coding sequence of *PpTIL* was replaced by the *NPTII* cassette using homologous recombination. Red arrows indicate positions of primers used for genomic PCR. **b,** Results of genomic PCR for four *PpTIL ko* lines using primers shown in **a**. The fragment length of genomic PCR is in line with expectation in **a**. **c,** Quantitative RT-PCR was performed to characterize four *PpTIL ko* mutants. *PpTIL* transcription level was evaluated for the wild type and four *PpTIL ko* lines using qRT-PCR. No expression of *PpTIL* was detected in the four *ko* mutants. Three biological replications were performed, and *PpEF1a* was used as reference gene for normalization. Data show means ± s.e.m.

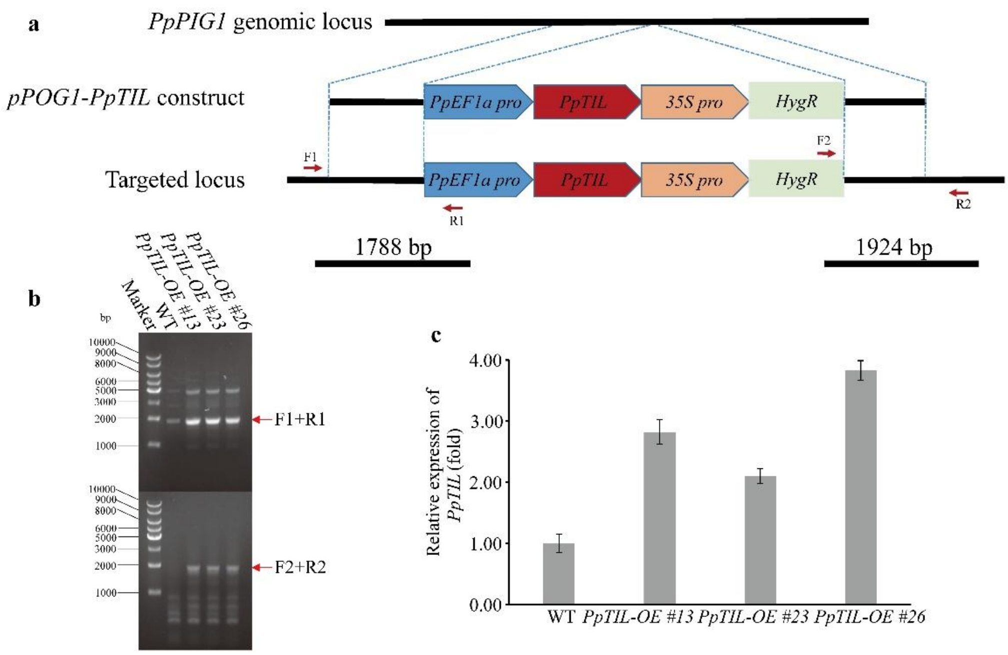


**Supplementary Fig. 5.** Generation and molecular identification of *PpTIL OE* lines. **a,** Schematic diagram of *PpTIL OE* construction and primers used for genotyping. Promoter *PpEF1a* was used for enhancing *PpTIL* expression in *PpPIG1* genomic locus, and hygromycin was used as selection marker and driven by 35S promoter. Red arrows denote positions of primers used for genomic PCR. **b,** Genomic PCR confirmation of three *PpTIL OE* lines using primers shown in **a**. The fragment length of genomic PCR is consistent with expectation. **c,** Quantitative RT-PCR was used to analyze three *PpTIL OE* lines. *PpTIL* transcription level was evaluated for the wild type and three *PpTIL OE* lines using qRT-PCR. The expression level of *PpTIL* is over two-fold higher in *PpTIL OE* lines than in the wild type. Three biological replications were performed and *PpEF1a* was used as reference gene for normalization. Data show means ± s.e.m.

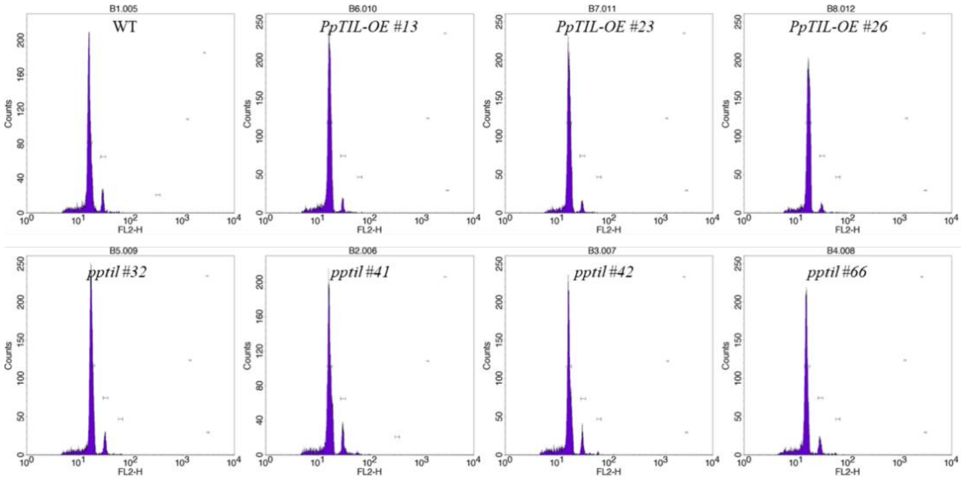


**Supplementary Fig. 6.** Chromosome ploidy analyses. WT, three *OE* and four *ko* plants were grown on BCDAT medium for one week, and their chromosome ploidy levels were analyzed using flow cytometry. The results indicate that chromosome ploidy levels of *ko* and *OE* lines are consistent with that of WT plants.

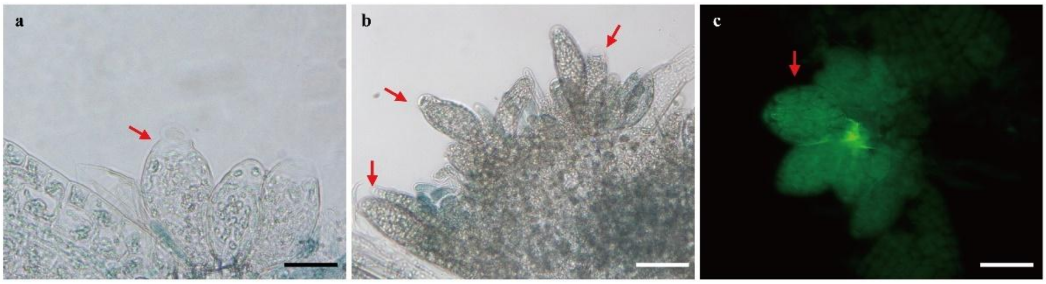


**Supplementary Fig. 7.** *PpTIL* is expressed in antheridia of *P. patens*. **a** and **b** show GUS signal in antheridia. **c** shows GFP signal in antheridia. Red arrows indicate antheridia bundles. Scale bars: 50 µm.

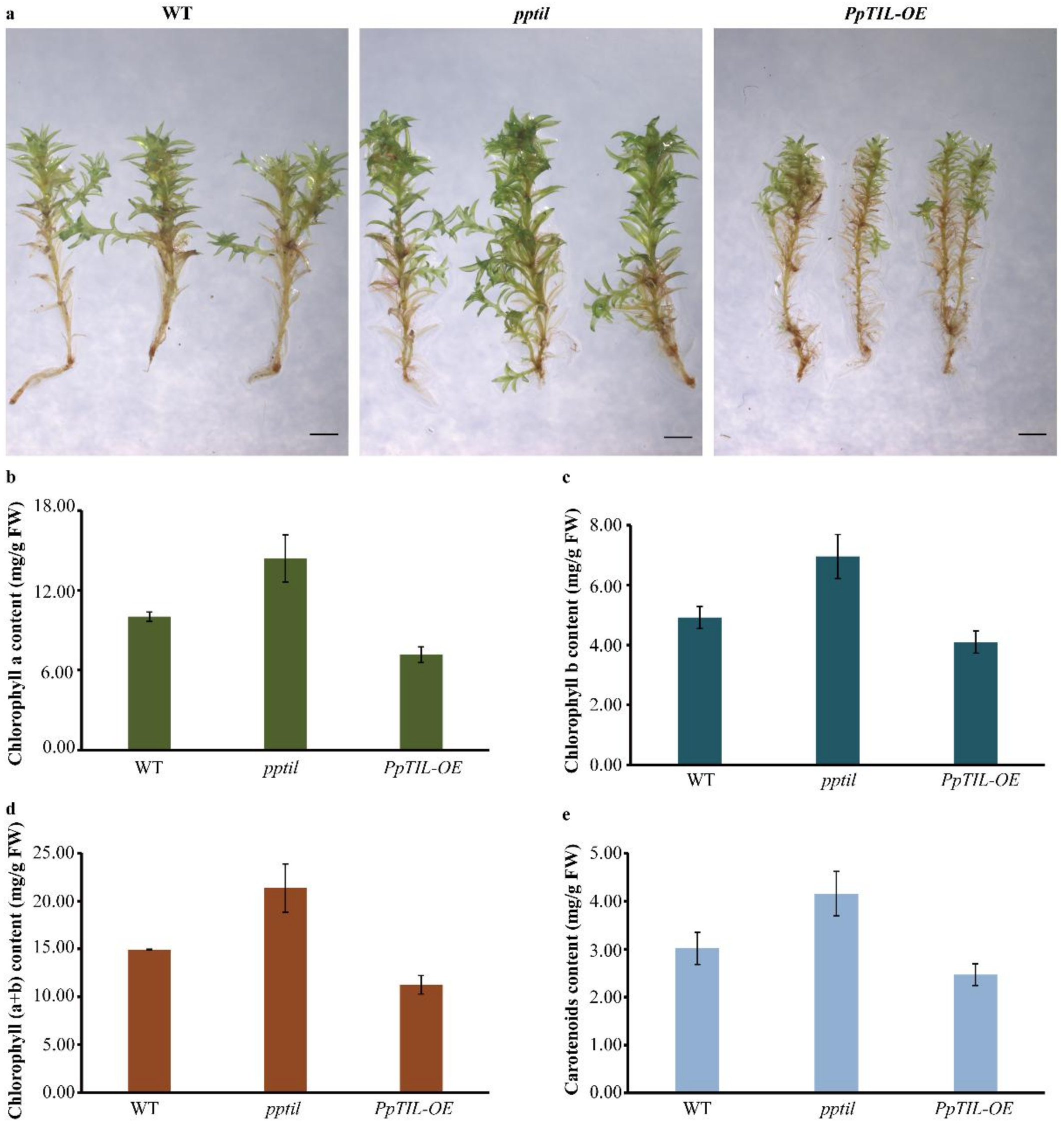


**Supplementary Fig. 8.** Phenotypes and pigment content of mature gametophores bearing archegonia and antheridia in WT, *ko* and *OE* plants. **a,** Phenotypes of mature gametophores of WT, *ko* and *OE* plants. Scale bar: 1 mm. **b-e,** The content of chlorophyll a, chlorophyll b, total chlorophyll (a+b), and carotenoids in WT, *ko* and *OE* plants. Three biological replications were performed. Data show means ± s.e.m.

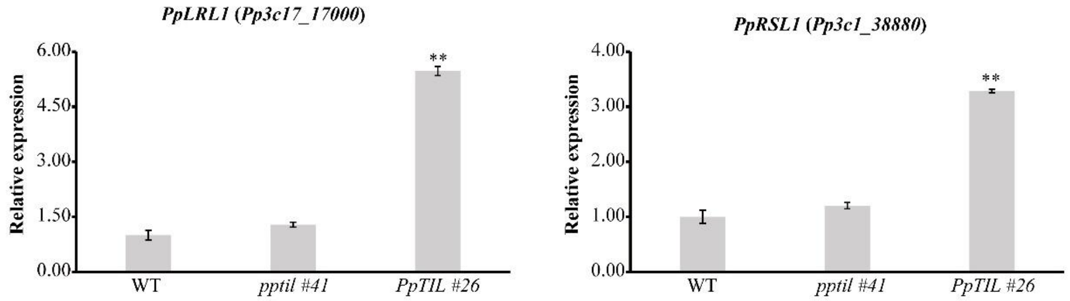


**Supplementary Fig. 9.** Relative expression of *PpLRL1* and *PpRSL1*. Quantitative RT-PCR was used to detect the expression level of *PpLRL1* and *PpRSL1* for WT, *PpTIL ko* and *OE* plants. Three biological replications were performed, and *PpEF1a* was used as reference gene for normalization. Data show means ± s.e.m. Asterisks indicate a statistically significant difference compared with the wild type based on a two-tailed Student's *t* test (** $p$<0.01).

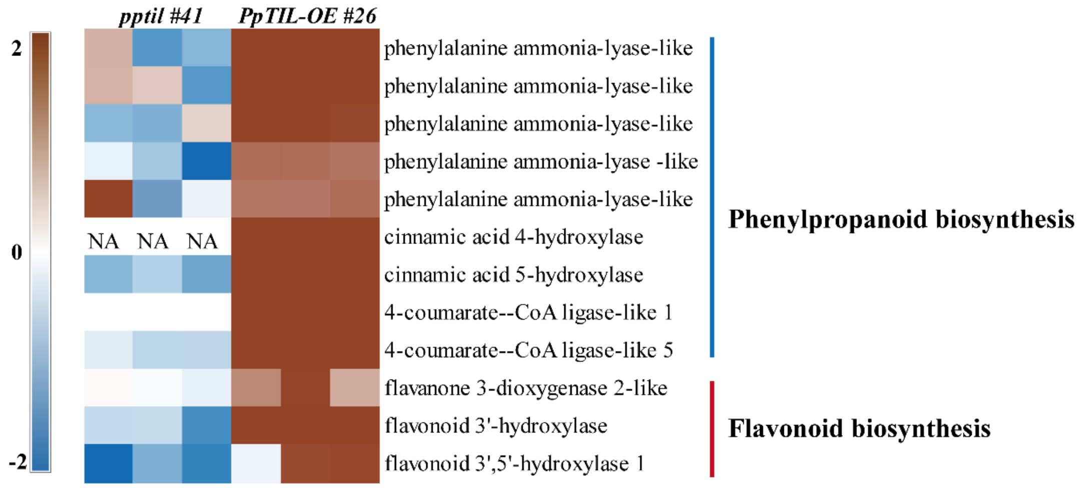


**Supplementary Fig. 10.** Genes involved in phenylpropanoid and flavonoid biosynthesis are differentially expressed in *ko* and *OE* plants of *PpTIL*. Among the differentially expressed are five genes encoding PAL (*Pp3c2_32410*, *Pp3c24_13110*, *Pp3c10_21810*, *Pp3c1_18940* and *Pp3c19_13690*), two genes encoding C4H (*Pp3c13_14870* and *Pp3c3_17840*), two genes encoding 4CL (*Pp3c8_730* and *Pp3c22_9051*), and three genes involved in flavonoid biosynthesis (*Pp3c27_4790*, *Pp3c2_2400,* and *Pp3c1_23870*). Additional information for each gene is provided in Supplementary Table 2.

a

| Gene ID | log2FoldChange | | | | | | Gene description |
|---|---|---|---|---|---|---|---|
| | *pptil #41* vs WT | | | *PpTIL-OE #26* vs WT | | | |
| Pp3c17_16260 | 0.68 | 0.53 | 0.31 | 1.94 | 1.95 | 1.91 | linoleate 9S-lipoxygenase-like |
| Pp3c7_25640 | -0.35 | -0.30 | -0.53 | 2.83 | 2.71 | 2.72 | allene oxide synthase-like |
| Pp3c5_3730 | -0.18 | 0.34 | 0.25 | 2.87 | 2.71 | 2.87 | allene oxide cyclase |
| Pp3c2_24500 | -3.57 | 0.90 | 1.55 | 2.78 | 3.19 | 2.85 | allene oxide cyclase |
| Pp3c4_22490 | 0.59 | 0.60 | 0.44 | 1.52 | 1.44 | 1.45 | allene oxide cyclase |

b

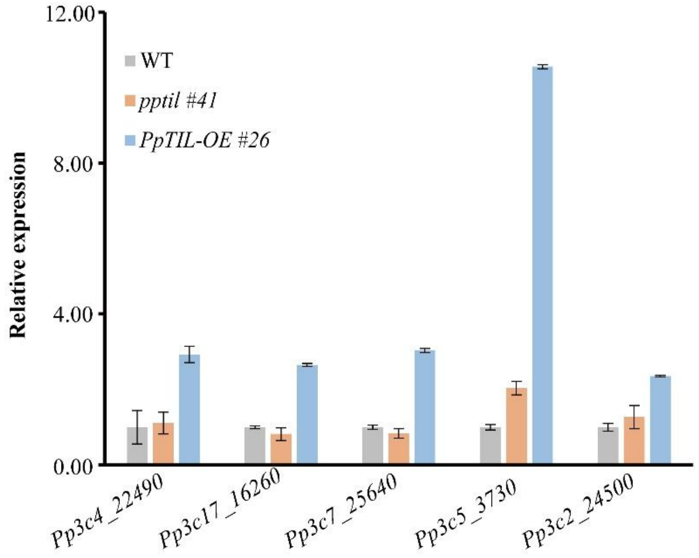


**Supplementary Fig. 11.** The expression of *LOX*, *AOS* and *AOC* genes in the OPDA biosynthetic pathway. **a,** Genes related to OPDA synthesis are up-regulated in *OE* mutants. Data were generated from RNA-seq data. **b,** The expression of *LOX*, *AOS* and *AOC* was verified through qRT-PCR. Three biological replications were performed, and *PpEF1a* was used as reference gene for normalization. Data show means ± s.e.m.

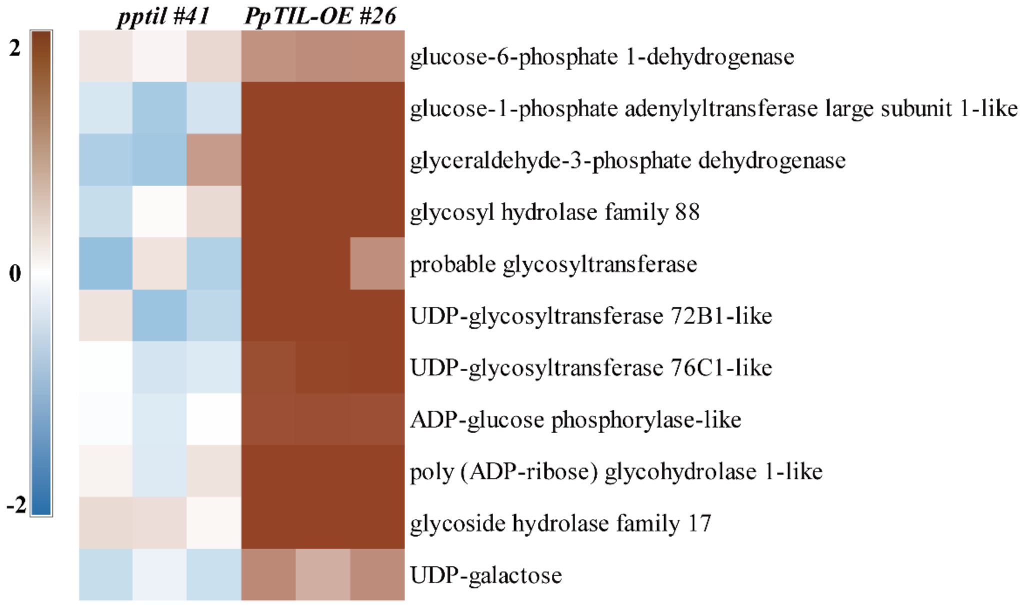


**Supplementary Fig. 12.** Genes involved in carbohydrate metabolism are differentially expressed in the *ko* and *OE* plants of *PpTIL*. *Pp3c23_16910*: glucose-6-phosphate 1-dehydrogenase; *Pp3c1_31540*: glucose-1-phosphate adenylytransferase large subunit 1-like; *Pp3c23_20200*: glyceraldehyde-3-phosphate dehydrogenase; *Pp3c5_25040* and *Pp3c16_1668*: glycosyl hydrolase family 88 and family 17; *Pp3c2_6500*: probable glycosyltransferase; *Pp3c12_18470* and *Pp3c16_5590*: UDP-glycosyltransferase 72B1-like and 76C1-like, respectively; *Pp3c17_20460*: ADP-glucose phosphorylase-like; *Pp3c3_12290*: poly (ADP-ribose) glycohydrolase 1-like; *Pp3c14_24680*: UDP-galactose.

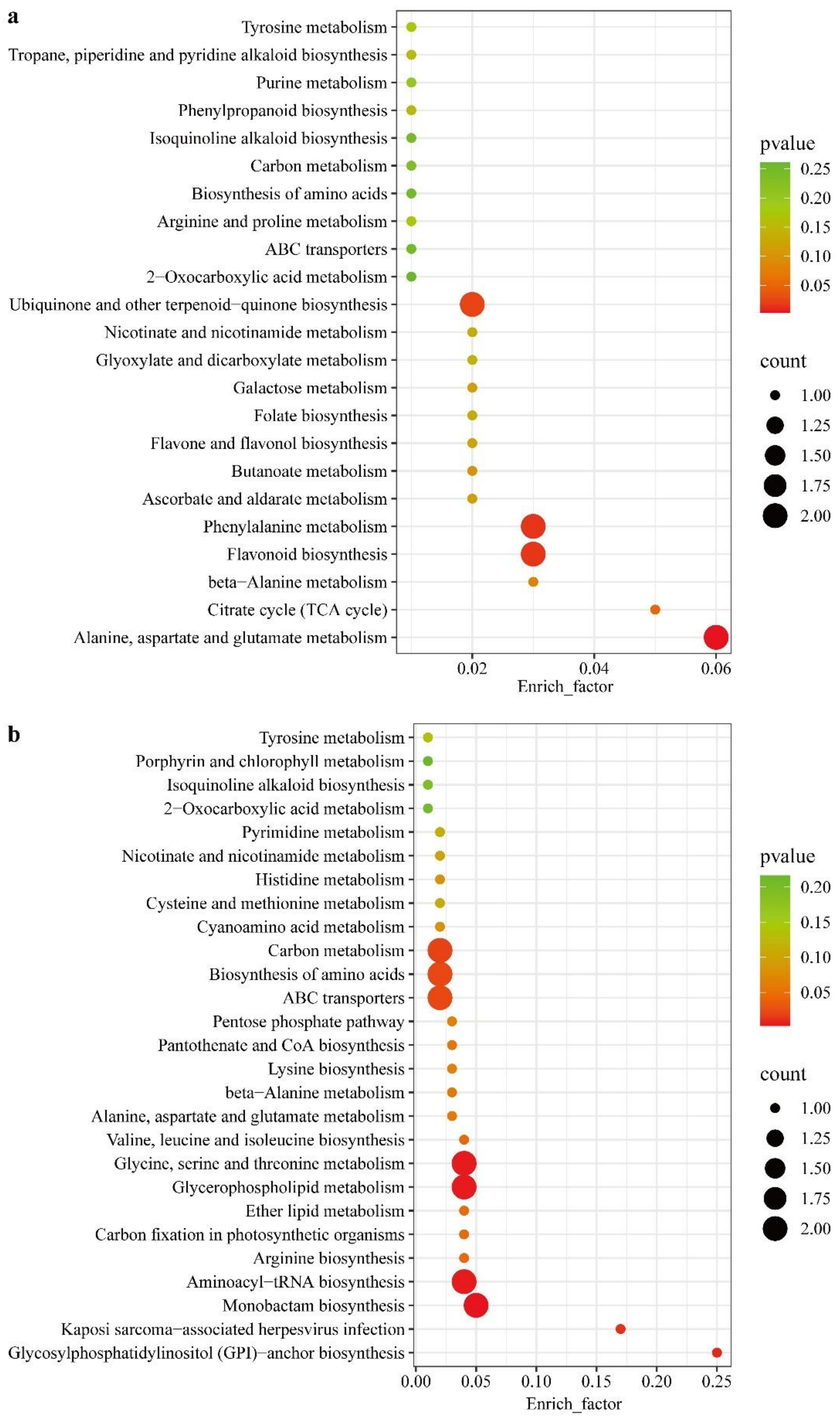


**Supplementary Fig. 13.** Results of KEGG enrichment analyses for differentially expressed metabolites in *ko* (**a**) and *OE* (**b**) plants relative to the wild type.

**Supplementary Table 1.** Additional gene information for Fig. 5f of the main text. These genes are related to auxin biosynthesis, transport, binding and response. Data were generated from RNA-seq data and verified through qRT-PCR.

| Gene ID | log2FoldChange | | | | | | Gene description |
|---|---|---|---|---|---|---|---|
| | *pptil #41* | | | *PpTIL-OE #26* | | | |
| Pp3c21_16440 | -0.48 | -0.42 | -0.55 | 1.32 | 1.31 | 1.36 | PpSHI1 |
| Pp3c23_10200 | 3.36 | 2.62 | 2.89 | -5.05 | -1.96 | -2.79 | PINA |
| Pp3c8_15410 | 0.59 | -0.16 | 0.00 | 4.36 | 4.37 | 4.31 | protein NRT1/ PTR family |
| Pp3c1_36410 | 0.46 | 0.72 | 0.42 | 4.08 | 4.01 | 3.97 | auxin transport |
| Pp3c9_3590 | -3.57 | 1.29 | -3.62 | 5.06 | 3.66 | 3.67 | auxin transporter-like protein |
| Pp3c25_12120 | 0.17 | 0.20 | -0.08 | 1.12 | 1.26 | 1.06 | auxin responsive protein |
| Pp3c17_6090 | -0.50 | -0.37 | -0.09 | 0.98 | 1.02 | 1.03 | auxin-binding protein |
| Pp3c25_12110 | 0.67 | 0.65 | 0.66 | 0.97 | 1.28 | 1.12 | auxin-responsive protein SAUR72-like |
| Pp3c6_26890 | 0.41 | 0.07 | 0.06 | 0.93 | 0.92 | 0.81 | auxin response factor |
| Pp3c24_16260 | 0.30 | 0.21 | -0.09 | 0.80 | 0.72 | 0.72 | GH3 auxin-responsive promoter |
| Pp3c23_5870 | 0.24 | 0.41 | 0.28 | 0.76 | 0.98 | 0.91 | protein auxin signaling F-BOX 2-like |
| Pp3c4_12750 | -3.18 | -2.76 | -2.26 | -0.90 | -0.35 | 1.51 | auxin-binding protein T85-like |
| Pp3c6_4560 | -1.49 | -1.39 | -1.56 | -2.19 | 0.23 | 2.31 | auxin-responsive protein SAUR66-like |

**Supplementary Table 2.** Additional gene information for Supplementary Fig. 10. These genes are related to phenylpropanoid and flavonoid biosynthesis. Data were generated from RNA-seq data.

| **Gene ID** | **log2FoldChange** | | | | | | **Gene description** |
|---|---|---|---|---|---|---|---|
| | ***pptil #41*** | | | ***PpTIL-OE #26*** | | | |
| Pp3c2_32410 | 0.82 | -1.32 | -0.93 | 2.94 | 2.98 | 3.00 | phenylalanine ammonia-lyase-like |
| Pp3c24_13110 | 0.81 | 0.58 | -1.31 | 3.16 | 3.00 | 3.00 | phenylalanine ammonia-lyase-like |
| Pp3c10_21810 | -0.89 | -0.99 | 0.48 | 2.06 | 2.09 | 1.97 | phenylalanine ammonia-lyase-like |
| Pp3c1_18940 | -0.16 | -0.71 | -2.81 | 1.58 | 1.56 | 1.46 | phenylalanine ammonia-lyase -like |
| Pp3c19_13690 | 2.04 | -1.37 | -0.17 | 1.43 | 1.42 | 1.55 | phenylalanine ammonia-lyase-like |
| Pp3c13_14870 | NA | NA | NA | 5.06 | 5.51 | 5.91 | cinnamic acid 4-hydroxylase |
| Pp3c3_17840 | -0.92 | -0.56 | -1.14 | 4.45 | 4.55 | 4.45 | cinnamic acid 5-hydroxylase |
| Pp3c8_730 | NA | NA | NA | 5.34 | 6.16 | 5.82 | 4-coumarate--CoA ligase-like 1 |
| Pp3c22_9051 | -0.22 | -0.52 | -0.49 | 3.04 | 3.12 | 2.96 | 4-coumarate--CoA ligase-like 5 |
| Pp3c27_4790 | 0.06 | -0.05 | -0.18 | 1.24 | 1.99 | 0.87 | flavanone 3-dioxygenase 2-like |
| Pp3c2_24000 | -0.44 | -0.43 | -1.48 | 5.01 | 5.01 | 4.88 | flavonoid 3'-hydroxylase |
| Pp3c1_23870 | -2.37 | -1.01 | -1.63 | -0.13 | 1.94 | 1.97 | flavonoid 3',5'-hydroxylase 1 |

**Supplementary Table 3.** Additional gene information for Fig. 6d of the main text. These genes are related to AP2-type transcription factors. Data were generated from RNA-seq data and verified through qRT-PCR.

| **Gene ID** | **log2FoldChange** | | | | | | **Gene description** |
|---|---|---|---|---|---|---|---|
| | ***pptil #41*** | | | ***PpTIL-OE #26*** | | | |
| Pp3c11_10660 | NA | NA | NA | 9.21 | 9.03 | 8.92 | AP2 |
| Pp3c15_3620 | NA | NA | NA | 7.60 | 7.61 | 7.48 | AP2 |
| Pp3c9_4590 | NA | NA | NA | 7.29 | 6.88 | 6.88 | AP2 |
| Pp3c3_6830 | NA | NA | NA | 3.87 | 4.01 | 3.88 | AP2 |
| Pp3c16_12850 | 0.33 | 0.20 | 0.17 | 3.74 | 3.75 | 3.67 | AP2 |
| Pp3c10_20000 | -0.50 | 1.22 | -0.27 | 3.73 | 3.81 | 3.70 | AP2 |
| Pp3c25_12880 | -0.61 | -0.38 | -1.10 | 3.73 | 3.73 | 3.82 | TINY-like |
| Pp3c7_20170 | 0.54 | -0.33 | -0.52 | 3.20 | 3.27 | 3.27 | AP2 |
| Pp3c5_4660 | 0.70 | -0.15 | -0.14 | 3.12 | 3.15 | 2.91 | AP2 |
| Pp3c5_2900 | 0.19 | -0.22 | 0.26 | 3.01 | 3.19 | 3.23 | AP2 |
| Pp3c2_36690 | NA | NA | NA | 2.92 | 3.49 | 3.34 | AP2 |
| Pp3c5_880 | 0.03 | 0.43 | 0.37 | 2.90 | 2.44 | 2.49 | AP2 |
| Pp3c4_2530 | 0.43 | 1.11 | -0.05 | 2.52 | 2.68 | 2.46 | AP2 |
| Pp3c9_25570 | 0.72 | 0.48 | 0.08 | 1.59 | 1.49 | 1.55 | AP2 |
| Pp3c7_2300 | 0.62 | -0.75 | -1.72 | 1.56 | 1.54 | 1.61 | AP2 |
| Pp3c6_28370 | -0.14 | -0.02 | -1.31 | 1.52 | 1.54 | 1.58 | AP2 |
| Pp3c20_3770 | 0.16 | 0.20 | -0.18 | 1.50 | 1.51 | 1.53 | AP2 |
| Pp3c26_1890 | -0.33 | -0.61 | -0.51 | 1.40 | 1.37 | 1.30 | AP2 |
| Pp3c26_13260 | -0.90 | -0.63 | 0.12 | 1.20 | 1.46 | 1.15 | AP2 |
| Pp3c1_7800 | -0.36 | -0.38 | -0.51 | 1.18 | 1.23 | 1.20 | AP2 |
| Pp3c6_19350 | NA | NA | NA | 1.13 | 0.97 | 0.82 | AP2 |
| Pp3c18_1800 | -0.21 | -0.07 | -0.18 | 1.12 | 1.06 | 1.14 | AP2 |
| Pp3c17_10170 | -0.05 | -0.03 | -0.33 | 1.05 | 1.35 | 1.15 | AP2 |
| Pp3c11_14690 | -1.75 | -0.29 | -1.06 | 0.93 | 1.33 | 1.24 | AP2 |
| Pp3c3_13510 | -0.77 | -0.45 | -1.03 | 0.87 | 0.57 | 0.42 | AP2 |
| Pp3c6_26080 | -0.29 | -0.24 | -1.55 | -0.36 | 0.30 | 0.80 | AP2 |

**Supplementary Table 4.** Additional gene information for Fig. 7a of the main text. These genes are related to lipid metabolism. Data were generated from RNA-seq data of *PpTIL ko* and *OE* plants compared to the wild type.

| **Gene ID** | **log2FoldChange** | | | | | | **Gene description** |
|---|---|---|---|---|---|---|---|
| | ***pptil #41*** | | | ***PpTIL-OE #26*** | | | |
| Pp3c12_7930 | NA | NA | NA | 6.75 | 7.02 | 6.62 | Phospholipase A1 |
| Pp3c23_14820 | 0.23 | 0.29 | 0.13 | 1.64 | 1.91 | 1.68 | lipid phosphate phosphatase 2 |
| Pp3c24_2640 | 2.20 | 0.91 | -0.61 | 4.12 | 4.06 | 3.75 | lipid phosphate phosphatase 2-like |
| Pp3c14_10640 | -4.95 | -1.95 | -0.15 | 3.72 | 3.57 | 3.66 | Lipoxygenase |
| Pp3c12_14480 | 0.44 | 0.45 | 0.27 | 1.76 | 1.70 | 1.57 | Lipoxygenase |
| Pp3c10_17370 | -3.56 | -3.59 | -3.61 | 1.19 | 1.17 | 1.24 | GDSL esterase/lipase 6-like |
| Pp3c9_2920 | -3.19 | -3.21 | -3.23 | 3.06 | 3.03 | 2.98 | GDSL esterase/lipase 6-like |
| Pp3c16_1160 | 0.35 | 0.23 | -0.07 | 1.34 | 1.43 | 1.38 | aldehyde oxidase GLOX-like |
| Pp3c12_15000 | -0.73 | -0.69 | -0.80 | 1.60 | 1.62 | 1.62 | aldehyde oxidase GLOX-like |

**Supplementary Table 5.** Additional information for Fig. 7b of the main text. Multiple pathways identified in KEGG enrichment analyses of DEGs are related to metabolism of lipids and their derivatives.

| **KEGG term** | **GeneRatio** | **P-value** | **padj** |
|---|---|---|---|
| Phenylpropanoid biosynthesis | 31/577 | 0.00000 | 0.00007 |
| Starch and sucrose metabolism | 34/577 | 0.00002 | 0.00121 |
| Plant hormone signal transduction | 26/577 | 0.00005 | 0.00192 |
| Circadian rhythm - plant | 19/577 | 0.00020 | 0.00527 |
| Phenylalanine metabolism | 13/577 | 0.00031 | 0.00557 |
| ABC transporters | 9/577 | 0.00031 | 0.00557 |
| alpha-Linolenic acid metabolism | 14/577 | 0.00132 | 0.02011 |
| Glycerophospholipid metabolism | 23/577 | 0.00170 | 0.02276 |
| Glycerolipid metabolism | 17/577 | 0.00198 | 0.02359 |
| MAPK signaling pathway - plant | 23/577 | 0.00252 | 0.02639 |
| Brassinosteroid biosynthesis | 7/577 | 0.00291 | 0.02639 |
| Flavonoid biosynthesis | 12/577 | 0.00296 | 0.02639 |
| Cysteine and methionine metabolism | 22/577 | 0.00382 | 0.03145 |
| Cutin, suberine and wax biosynthesis | 6/577 | 0.00718 | 0.05250 |
| Nitrogen metabolism | 11/577 | 0.00737 | 0.05250 |
| Glutathione metabolism | 16/577 | 0.00785 | 0.05250 |
| Ether lipid metabolism | 10/577 | 0.01217 | 0.07392 |
| Isoquinoline alkaloid biosynthesis | 8/577 | 0.01244 | 0.07392 |
| Folate biosynthesis | 9/577 | 0.01672 | 0.09415 |
| Pentose and glucuronate interconversions | 15/577 | 0.01777 | 0.09505 |

**Supplementary Table 6.** Primers used for vector construction of *KI*, *ko* and *OE* of *PpTIL*.

| **Primer** | **Sequence (5'-3')** |
|---|---|
| pTN85-PpTIL-KI-5F(KpnI) | GAACAAAAGCTGGGTACCTTGAATGGTGCTAACGAGTC |
| pTN85- PpTIL-KI-5R(XhoI) | GCTCACGTCGACCTCGAGCTTTCCGAAGAGGGATTTT |
| pTN85-PpTIL-KI-3F(BamHI) | CGGGGATCGGGGGGATCCAAACCAATGATGCACATGGC |
| pTN85-PpTIL-KI-3R(XbaI) | GGTGGCGGCCGCTCTAGAGTTGTGTCATTGAGAGAAGCA |
| pTN182-PpTIL-ko-5F(KpnI) | TAGGGCGAATTGGGTACC AGGGCAAAGAGACAGCAGCA |
| pTN182-PpTIL-ko-5R(SalI) | CTTATCGATACCGTCGACCAGACGAGGGTATCAACGAC |
| pTN182-PpTIL-ko-3F(XbaI) | CGCATGCCCGGGTCTAGA TCTTTCGTGAGTTTTGCCCT |
| pTN182-PpTIL-ko-3R(BmaHI) | CGGCCGCTCTAGGATCC GTTGTGTCATTGAGAGAAGCA |
| pPOG1-PpTIL-OE-F(NotI) | TATCCAGTCACTATGGCGGCCGCATGGGAGGCGAAAAGGACTT |
| pPOG1-PpTIL-OE-F(XhoI) | TTCTCCTTTACCCATCTCGAGCTTTCCGAAGAGGGATTTTAAC |

**Supplementary Table 7.** Primers used for genotyping of *KI*, *ko* and *OE* plants of *PpTIL*.

| **Primer** | **Sequence (5'-3')** |
|---|---|
| PpTIL-KI-F1 | TTTCCCAAGACACACTTACAC |
| PpTIL-KI-R1 | AGTTCACCTTGATGCCGTTC |
| PpTIL-KI-F2 | TAAACCAGACACGAGACGAC |
| PpTIL-KI-R2 | GATGCCTATCAAAGAGATCAAC |
| PpTIL-ko-F1 | TTTCCCAAGACACACTTACAC |
| PpTIL-ko-R1 | ATAGTGGGATTGTGCGTCAT |
| PpTIL-ko-F2 | TAAACCAGACACGAGACGAC |
| PpTIL-ko-R2 | GATGCCTATCAAAGAGATCAAC |
| PpTIL-OE-F1 | TTTCCTAAGTCGGGTCCTGT |
| PpTIL-OE-R1 | TCCATCTGTCCTAACTATCA |
| PpTIL-OE-F2 | TTGACTCCCCCGTAGGTTTG |
| PpTIL-OE-R2 | CCAGATTAGCAAAGCCACCC |

**Supplementary Table 8.** Primers used for qRT-PCR of *PpTIL* and genes related to auxin, AP2 domain, OPDA biosynthesis, *PpLRL1*, *PpRSL1* and *PpEF1α*.

| Gene ID | Primer-F (5'-3') | Primer-R (5'-3') |
|---|---|---|
| PpTIL | CAGCCCTGACGCTAAACTCA | GCTCTGGCGTCCGAGATAAA |
| Pp3c21_16440 | CCGCGGCTCTGATATCAACT | CATCTTCCACGCCTGTCACT |
| Pp3c23_10200 | AAACGTCAACGCAACTGCTC | AAACCAGCTTCAGAGACGGG |
| Pp3c8_15410 | GGCAAGCTCAACACAACTGG | TGTACACCAACGCAGAGGAC |
| Pp3c1_36410 | ATGGAACAAGGAGGCCGAAG | ATTGGAGGTGCTATACGGCG |
| Pp3c9_3590 | AAGGAGAGAGAGGGCCACAA | TCACAGCTCCAAACAGGCAA |
| Pp3c25_12110 | TGATAGCAACGACTCTGCCC | CTAACCCTCCCTTGTGCTCG |
| Pp3c17_6090 | GTTTGTGGCTCCTCGTGAGA | AGTTCATGTAGGGCGTGTGG |
| Pp3c25_12120 | AGAGTTCGGGTTCTGTTGCG | AACTCTGCAGTAGCACCCTC |
| Pp3c6_26890 | TGCCTCATGTGGGTGCTAAG | ATCTCAGGCTGCAGGCAAAT |
| Pp3c24_16260 | GGGAGATGGTGGATAGCAGC | CCATCACGCGATTGAACGTC |
| Pp3c23_5870 | GAGATTGTCCCTTCGGCGAT | TGAGCGATGGATTGTTGGCT |
| Pp3c4_12750 | GCATCAGGTGAGGAACACGA | TCCCACTCTCTTGGCTCGTA |
| Pp3c6_4560 | GCCTTTCGGGCGTTATTAGC | AACAATCCTGTCCCACGACC |
| Pp3c11_10660 | GGCCAATCTGAACTTCCCCA | GCTCTTCGTCACTGCTGTCT |
| Pp3c15_3620 | TATCGCGGTGTTCGGATGAG | TGGATTTCGCGTGTGGTGTA |
| Pp3c9_4590 | ATTTCATGTCGCCTGGCTCA | TTCCTCGAGCCATCTCATGC |
| Pp3c3_6830 | GTTGTCTCTGCCAAATCCGC | ATGCAGACGAACGCGATACA |
| Pp3c16_12850 | AGCTTGTCGCAGAGACTGAC | ATCGACAAACACTACCGCTG |
| Pp3c10_20000 | CCAGTACTCGCAACCGAACT | ATCCCGCTCAAAGTGGACTG |
| Pp3c25_12880 | ACTATCTGGCTGGGCACCTA | TTCAGCATTGTCTAGCCGCA |
| Pp3c7_20170 | CTCTTCACTCCGCAACCCTT | TGCAATAACCTGCTTCCGGT |
| Pp3c5_4660 | ATCAGCTCAAGAGACGCCTGC | TCCACGCGTTCGCTAGAATC |
| Pp3c5_2900 | GCGTGCCCAGATCTACTACC | GTATCGTATGTGCCGAGCCA |
| Pp3c2_36690 | TGGTAGGCGGTTTTCGTTCA | TACTGCGATGACGACGTTCC |
| Pp3c5_880 | CGGAATCATACCTCCCACCG | AGGGGCTGGTTGGGTACTAT |
| Pp3c4_2530 | AAAGATCCCGAGCACACGTT | GGTCTGCGGTAGAGGATTCG |
| Pp3c9_25570 | TCTGTGTTCAGAGGCGTCAC | TAGGCTCTGGCTGCCTTTTC |
| Pp3c7_2300 | ACGTCCTCCGAGACACCTTA | TTGTGCATGCCGATGGTTTG |
| Pp3c6_28370 | ACCGAGAGACCTGGAACGTA | GGTGGTGTGCTTAAGTCGGA |
| Pp3c20_3770 | GCCTCCGTTACCAGAGATCG | TCCGAGGTACTCCAATCCGT |
| Pp3c26_1890 | CCTCAAAGTCTCGTCCACCC | ACTAGTGGAGCTGGTCGTCT |

| | | |
|---|---|---|
| Pp3c26_13260 | GCCATCCTCCTCTCCGAAAC | ATGCGGTAATTCCGTCCCAG |
| Pp3c1_7800 | CCGACATTCCAGCTCCTGTT | GCTTTCGACGATTCCGTTCG |
| Pp3c6_19350 | CAATCCAGTAACCGCCCAGT | TGTTGGCGCACCATCAATTC |
| Pp3c18_1800 | GCAAGGTCAAAGAACAGGCG | GCTAGTGTCTTTCGCCGCTA |
| Pp3c17_10170 | AGCGTCCTTCTTCAACTCCG | TCAACTTGCGAGACCGAGAC |
| Pp3c11_14690 | ACTCACCATTGCCGTCTCTG | CCGACCCAATTCTGGACGAT |
| Pp3c3_13510 | TCTGGGCCCTAACTTTGCAG | CGGTAGGACTCGAAGATGGC |
| Pp3c6_26080 | GATGCTACGCCGTACTCGAA | TTCTGATCGACGCTACTCGC |
| Pp3c17_16260 | TCCCTAGCAGGGAGAAGGAC | CTCAACTGCCTGTTGGTTGC |
| Pp3c7_25640 | TCCCCTACTTCGGTGCCATA | TGACACGAAACACCGTGCTA |
| Pp3c5_3730 | TTTGCAGCCCGCAATCTCTA | GGCAAAAACGCAGGACTGTT |
| Pp3c2_24500 | TTGAGAGCACCTTGCTTGCC | GGACGGATCTGATGGAGGAC |
| Pp3c4_22490 | TCGGAGGAAAGAAACAGCCG | TGAGAGATCAGCGTGCAGAG |
| Pp3c1_38880 | CTTGATGCATCTGCCGAAGC | TCATCACGGATGCACTGCTT |
| Pp3c17_17000 | GGTTTGGAGTGGGAGGGATG | TGATCGCGTCAGTGTATCCG |
| PpEF1α | AATCATACATTTCACCTCGCC | GATCAGTGGGTAGAAGTGAC |

**Supplementary Table 9.** Primers used for vector construction of luciferase complementation imaging (LCI) analysis.

| Primer | Sequence (5'-3') |
|---|---|
| PpTIL-nLUC-KpnI-F | GACGAGCTCGGTACCATGGGAGGCGAAAAGGACTT |
| PpTIL-nLUC-SalI-R | CGAGATCTGGTCGACCTTTCCGAAGAGGGATTTTAAC |
| Pp3c2_24110-cLUC-KpnI-F | TCCCGGGGCGGTACCATGGCGGATCAGAACTCTGC |
| Pp3c2_24110-cLUC -SalI-R | GCTCTGCAGGTCGACTCATGCAAGGCCTGCATCCT |
| Pp3c21_21000-cLUC-KpnI-F | TCCCGGGGCGGTACCATGGCCAGATTGTCCTTGTG |
| Pp3c21_21000-cLUC -SalI-R | GCTCTGCAGGTCGACTTATGTCCTGGCGTCCCAGA |
| Pp3c6_7260-cLUC-KpnI-F | TCCCGGGGCGGTACCATGATGGGAGGCGGCAAAGA |
| Pp3c6_7260-cLUC -SalI-R | GCTCTGCAGGTCGACTTAATGCCGTTGTTGCAACC |
| Pp3c3_8370-cLUC-KpnI-F | TCCCGGGGCGGTACCATGGTTGGGCTCACCATGG |
| Pp3c3_8370-cLUC -SalI-R | GCTCTGCAGGTCGACTCACTTCGTTTTGTTGACCC |
| Pp3c23_14000-cLUC-KpnI-F | TCCCGGGGCGGTACCATGCAACGTTTGCGAAAAGC |
| Pp3c23_14000-cLUC -SalI-R | GCTCTGCAGGTCGACTTACTTCTTCAACAGAAGGGG |
| Pp3c5_3730-cLUC-KpnI-F | TCCCGGGGCGGTACCATGGCGATGGCTGTGACCAA |
| Pp3c5_3730-cLUC -SalI-R | GCTCTGCAGGTCGACTTAGTCAGTGTAGTTGGGGA |
| Pp3c7_14500-cLUC-KpnI-F | TCCCGGGGCGGTACCATGGGCGTGGAGCAGGGGTTAA |
| Pp3c7_14500-cLUC -SalI-R | GCTCTGCAGGTCGACTTATGCTGAAGCCAGCGAACC |
| Pp3c3_25360-cLUC-KpnI-F | TCCCGGGGCGGTACCATGGCGGAAGAGACTATGGA |
| Pp3c3_25360-cLUC -SalI-R | GCTCTGCAGGTCGACCTAGTTTCGGTGACCCATGC |
| Pp3c5_20840-cLUC-KpnI-F | TCCCGGGGCGGTACCATGGGACCTGCCCAAGAGCCAGAT |
| Pp3c5_20840-cLUC -SalI-R | GCTCTGCAGGTCGACTCATTGCTTCTTCTGGCTAGCGTAG |
| Pp3c1_12020-cLUC-KpnI-F | TCCCGGGGCGGTACCATGCCTCCTGCAGTGCATTC |
| Pp3c1_12020-cLUC -SalI-R | GCTCTGCAGGTCGACTTACTTGTTTCGGAGACTTATGAGC |
| Pp3c14_19130-cLUC-KpnI-F | TCCCGGGGCGGTACCATGATCGCGATGGATAATTCG |
| Pp3c14_19130-cLUC -SalI-R | GCTCTGCAGGTCGACCTACGACAGAAACACATGCG |
| Pp3c17_770-cLUC-KpnI-F | TCCCGGGGCGGTACCATGAGCGCTCCGGCGAAAAA |
| Pp3c17_770-cLUC -SalI-R | GCTCTGCAGGTCGACCTAAGATGACTGAGACACAGGC |
| Pp3c2_5450-cLUC-KpnI-F | TCCCGGGGCGGTACCATGGGTCTGGACGAGGATTT |
| Pp3c2_5450-cLUC -SalI-R | GCTCTGCAGGTCGACTCAAGCCTCCTGGAGCTGCT |
| Pp3c15_14720-cLUC-KpnI-F | TCCCGGGGCGGTACCATGGCGCCTTACTCGGGTAA |
| Pp3c15_14720-cLUC -SalI-R | GCTCTGCAGGTCGACCTAGTACACGTAGTTCTTCACATG |
| Pp3c16_18700-cLUC-KpnI-F | TCCCGGGGCGGTACCATGGCGTTGGTCAAAGCGAA |
| Pp3c16_18700-cLUC -SalI-R | GCTCTGCAGGTCGACTTATTCGATTTCCTTGATTTGGGGC |

| | |
|---|---|
| Pp3c1_5050-cLUC-KpnI-F | TCCCGGGGCGGTACCATGGCTGCTAATGCTATGCT |
| Pp3c1_5050-cLUC -SalI-R | GCTCTGCAGGTCGACTCACTTCCAGAGCTCCTCCA |
| Pp3c3_32110-cLUC-KpnI-F | TCCCGGGGCGGTACCATGACGGTCACCGATTGTCGA |
| Pp3c3_32110-cLUC -SalI-R | GCTCTGCAGGTCGACCTACTCACTCAGGCCGCCACGTT |
| Pp3c10_10490-cLUC-KpnI-F | TCCCGGGGCGGTACCATGGAGGTTCTTCCCCGCGCTA |
| Pp3c10_10490-cLUC -SalI-R | GCTCTGCAGGTCGACTCAAACGGATTTCGCTGCAGAAG |
| Pp3c9_17550-cLUC-KpnI-F | TCCCGGGGCGGTACCATGGGGGGGATAAGGGATTT |
| Pp3c9_17550-cLUC -SalI-R | GCTCTGCAGGTCGACCTAAGATCTATCCGCCCGG |
| Pp3c7_24650-cLUC-KpnI-F | TCCCGGGGCGGTACCATGGCTGGAGCAGTGATGG |
| Pp3c7_24650-cLUC -SalI-R | GCTCTGCAGGTCGACCTAAAAGTCTAGACCAGCTTCAC |
| Pp3c10_18400-cLUC-KpnI-F | TCCCGGGGCGGTACCATGCCAACCTCTGTAGCGG |
| Pp3c10_18400-cLUC -SalI-R | GCTCTGCAGGTCGACTCACTTCAAGTTGGGAAGAT |
| Pp3c22_7570-cLUC-KpnI-F | TCCCGGGGCGGTACCATGGCAGCCCAGGTCGGAA |
| Pp3c22_7570-cLUC -SalI-R | GCTCTGCAGGTCGACCTATGCGAGCACGGCGAGGAT |
| Pp3c15_17530-cLUC-KpnI-F | TCCCGGGGCGGTACCATGGCTATGGCGCTGCAATC |
| Pp3c15_17530-cLUC -SalI-R | GCTCTGCAGGTCGACCTAGTCTTTCTTGTCCTCTGCC |
| Pp3c24_17080-cLUC-KpnI-F | TCCCGGGGCGGTACCATGGCGATGGCAGTGGGGA |
| Pp3c24_17080-cLUC -SalI-R | GCTCTGCAGGTCGACCTAGGCTGCTGTGGGTTCTGA |
| Pp3c4_31020-cLUC-KpnI-F | TCCCGGGGCGGTACCATGGCTTCCTGCGCTCTGCT |
| Pp3c4_31020-cLUC -SalI-R | GCTCTGCAGGTCGACTCAGTCCAAAACCTCGAATTCGT |
| Pp3c23_7670-cLUC-KpnI-F | TCCCGGGGCGGTACCATGGCCGCTCTCGCCA |
| Pp3c23_7670-cLUC -SalI-R | GCTCTGCAGGTCGACCTAAGCGCTGTCACCTCGTT |
| Pp3c24_4980-cLUC-KpnI-F | TCCCGGGGCGGTACCATGTTGCCATCATCTCAAGC |
| Pp3c24_4980-cLUC -SalI-R | GCTCTGCAGGTCGACCTAAGCAGCTGCAAGAGAAG |
